# An intranuclear bacterial parasite of deep-sea mussels expresses apoptosis inhibitors acquired from its host

**DOI:** 10.1101/2023.06.11.544495

**Authors:** Miguel Ángel González Porras, Adrien Assié, Målin Tietjen, Marlene Violette, Manuel Kleiner, Harald Gruber-Vodicka, Nicole Dubilier, Nikolaus Leisch

**Affiliations:** Max Planck Institute for Marine Microbiology, Bremen, Germany; Alkek Center for Metagenomics and Microbiome Research, Baylor College of Medicine, Houston, TX 77030, USA; Department of Plant and Microbial Biology, North Carolina State University, Raleigh, NC 27695, USA

**Keywords:** Laser-capture microdissection, lateral gene transfer, ankyrin repeats, *Endozoicomonas*, bathymodioline mussels, *Bathymodiolus*, *Gigantidas*, herpes virus, OsHV-1

## Abstract

Endozoicomonadaceae bacteria are widespread in many marine animals, and generally considered beneficial. Members of one clade, however, Candidatus Endonucleobacter, infect the nuclei of deep-sea mussels, where they replicate to ≥ 80,000 bacteria per nucleus and cause the nuclei to swell to 50 times their original size. How these parasites are able to persist in host nuclei without the cell undergoing apoptosis is not known. We show here that Ca. Endonucleobacter encodes and expresses 7-15 inhibitors of apoptosis (IAPs), proteins previously only known from animals and viruses. Dual RNA-seq transcriptomes of infected nuclei revealed parallel upregulation of Ca. Endonucleobacter IAPs and host caspases, suggesting an arms race between the parasite and host for control of apoptosis. Comparative phylogenetic analyses revealed that Ca. Endonucleobacter acquired IAPs repeatedly through horizontal gene transfer from their hosts in convergent acquisition, possibly mediated by herpes viruses that may infect both the parasite and the host.

## Introduction

Most metazoans are intimately associated with bacteria (McFall-Ngai et al., 2013), and some of these live inside eukaryotic cells or very rarely, inside eukaryotic organelles (Gruber-Vodicka et al., 2019; Sacchi et al., 2004; Salje, 2021). Marine animals are often associated with a family of gammaproteobacteria fittingly named Endozoicomonadaceae. Most Endozoicomonadaceae are extracellular, and only a few *Endozoicomonas* species and their close relatives live inside their host’s cells (Cano, 2020; Cano et al., 2018; Hooper et al., 2019; Howells et al., 2021). First isolated from a sea slug only 16 years ago (Kurahashi & Yokota, 2007), culture-independent sequencing approaches have revealed that *Endozoicomonas* and other Endozoicomonadaceae are ubiquotous and common inhabitants in a wide diversity of marine animals, from sponges and corals to fish (reviewed in Neave et al., 2016). Their role for their hosts has often been inferred but rarely proven, and is described as ranging from parasitic and commensalistic to beneficial (Katharios et al., 2015; Neave et al., 2016; Pogoreutz et al., 2022; Qi et al., 2018). All cultured Endozoicomonadaceae are aerobic, or facultatively anaerobic, heterotrophs that were isolated from marine hosts (Bartz et al., 2018).

A single clade of Endozoicomonadaceae, *Candidatus* Endonucleobacter, lives inside its host’s nuclei. These bacteria infect the nuclei of deep-sea bathymodioline mussels from hydrothermal vents and cold seeps around the world (Zielinski et al., 2009). The *Ca*. Endonucleobacter infection cycle begins with a single bacterium that invades the nucleus and then grows by elongating and dividing. In the final stages of infection, the elongated cells undergo septated division and replicate to as many as 80,000 cells, causing the mussel’s nuclei to swell to as much as 50 times their original size. Eventually, the infected mussel cells burst, releasing *Ca*. Endonucleobacter into the seawater (Zielinski et al., 2009).

Intranuclear bacteria have rarely been described in animals, but are well known from amoeba and other microbial eukaryotes (reviewed in Schulz & Horn, 2015). In microbial eukaryotes, the bacteria belong to other bacterial lineages than *Ca*. Endonucleobacter, such as the Rickettsiales and Verrucomicrobia, and do not replicate to such high numbers within their host’s nuclei as *Ca*. Endonucleobacter. To date, nothing is known about the molecular and cellular processes that intranuclear bacteria of animals use to infect and reproduce in their host. Key questions are how bacteria that infect animal nuclei are able to counter host immune responses, avoid the induction of host cell death through apoptosis, and how these parasites gain nutrition for their massive replication. Zielinski et al. (2009) hypothesized that *Ca*. Endonucleobacter digests nuclear chromatin but this would quickly impair the cellular activity of the host cell, including its immune responses (Schulz & Horn, 2015). Moreover, chromatin degradation, together with the deformation of the host cytoskeleton that *Ca*. Endonucleobacter induces through the dramatic increase in nuclear volume, would trigger apoptosis, a common response of metazoans to infection by parasites, and quickly lead to the death of infected cells (Ding et al., 2012; Niebuhr et al., 2000, 2002; Tavares et al., 2017).

To reveal the genetic adaptations that allow *Ca*. Endonucleobacter to thrive in its intranuclear niche, we assembled high quality genomes of two *Ca*. Endonucleobacter species, specific to two bathymodioline host species, *Bathymodiolus puteoserpentis* from hydrothermal vents on the Mid-Atlantic Ridge and *Gigantidas childressi* from cold seeps in the Gulf of Mexico, and compared them to closely related Endozoicomonadaceae. To gain insights into the metabolism of *Ca*. Endonucleobacter, we analyzed the metatranscriptomes and metaproteomes of bulk gill tissues from *G. childressi*. Finally, to understand host-microbe interactions during the infection cycle of *Ca*. Endonucleobacter, we used laser-capture microdissection, coupled with ultra-low-input dual RNA-seq, to generate transcriptomes of both the parasite and the host in early, middle and late infection stages (Fig. 1, a; Supplementary Information Video 1).

**Fig. 1.**
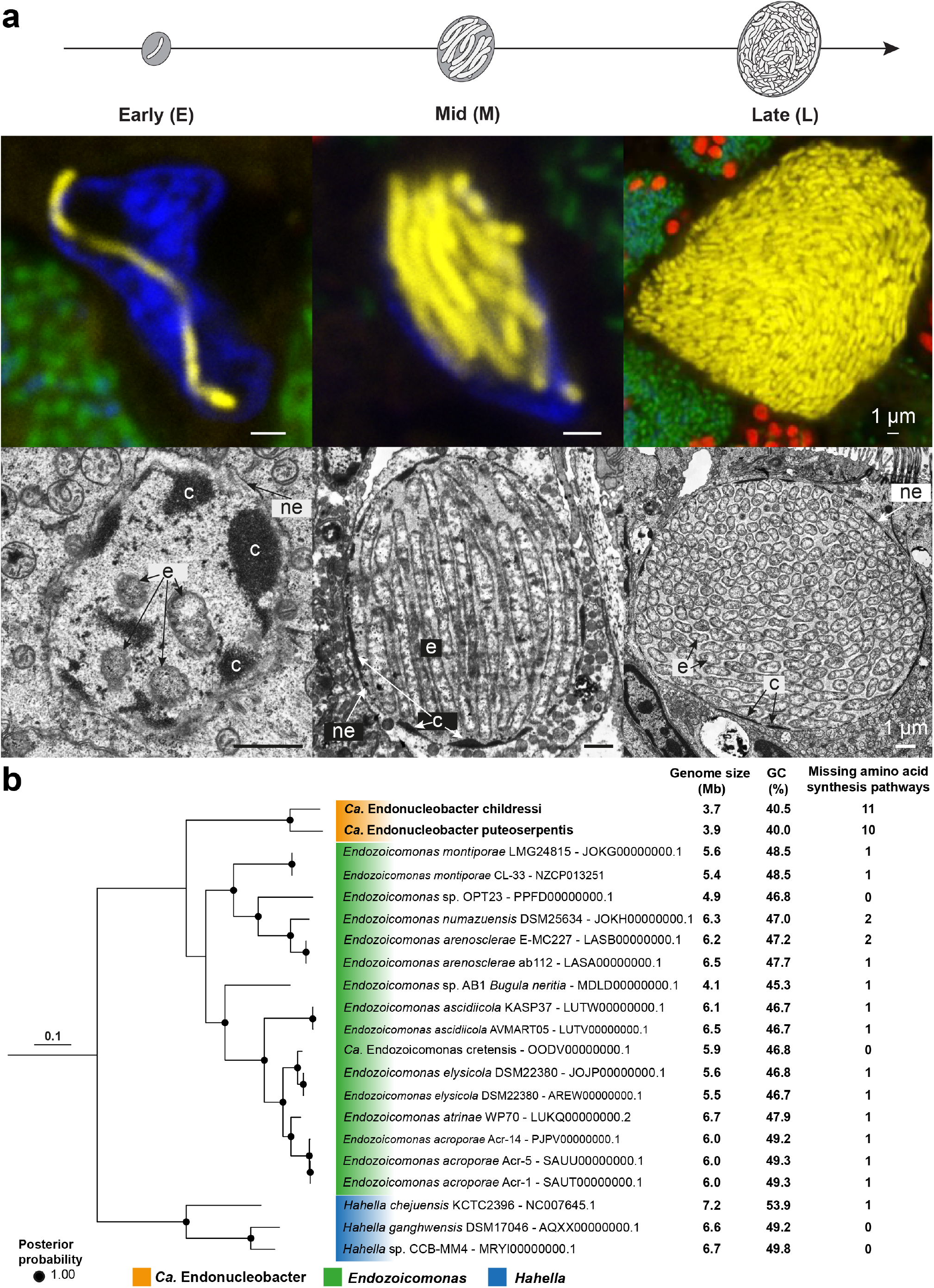
A single *Ca*. Endonucleobacter infects the mussel nucleus (early infection stage), then grows through elongation and division (mid infection stage), and finally divides through septation of the elongated cells to as many as 80,000 cells (late infection stage). In the final infection stage the nucleus is enlarged by as much as 50-fold in volume, the host cell bursts, and the parasites are released to the environment. (a), *Ca*. Endonucleobacter infectious cycle in early, mid and late stages of infection, shown respectively in the left, middle and right columns (diagram in top row, fluorescence *in situ* hybridization (FISH) in middle row, and transmission electron microscopy (TEM) in bottom row). FISH images show *Ca*. Endonucleobacter in yellow, the sulfur-oxidizing symbiont in green, the methane-oxidizing symbiont in red and DAPI-stained DNA in blue (sequences of all FISH probes are listed in Supplementary Information Table 2). Abbreviations: e: *Ca*. Endonucleobacter cell. c: chromatin. ne: nuclear envelope. (b), Phylogenomic analysis based on 514 single-copy genes shared between the genera *Ca*. Endonucleobacter and *Endozoicomonas*. Three *Hahella* genomes were used to root the tree. Key genome characteristics are listed to the right.

## Results and discussion

### Two *Ca*. Endonucleobacter species with different infection patterns

We used fluorescence *in situ* hybridization (FISH) analyses, with probes specific to *Ca*. Endonucleobacter, to analyze its distribution in *B. puteoserpentis* and *G. childressi*. These deep-sea mussels, like all other bathymodioline species investigated so far, house symbiotic sulfur- and/or methane-oxidizing bacteria in their gill cells that provide them with nutrition (Dubilier et al., 2008). Our FISH analyses confirmed that in both mussel host species, the parasite only infected the nuclei of gill cells that did not have sulfur- and methane-oxidizing symbionts, as previously reported (Zielinski et al., 2009). In *G. childressi, Ca*. Endonucleobacter was always restricted to the outer ciliated edges of the gill (**Extended Data Fig. 1a-c**), while in *B. puteoserpentis*, the parasite was distributed evenly across gill tissues (**Extended Data Fig. 1d-f**). The confinement of *Ca*. Endonucleobacter to the outer edges of the gill in *G. childressi* was fortunate because it allowed us to gain samples from these non-model, deep-sea hosts that were greatly enriched in the parasite, thus providing enough DNA for long-read sequencing and enabling the dual-seq RNA approaches described below.

Our analyses of high quality genomes, assembled from both short- and long-read sequencing of *B. puteoserpentis* and *G. childressi* gill tissues, revealed that these two mussel species are infected by genetically distinct *Ca*. Endonucleobacter species, based on their average nucleotide identity of 84.3%. The two *Ca*. Endonucleobacter species were named after their hosts, that is *Ca*. E. puteoserpentis and *Ca*. E. childressi (**Supplementary Note 1**). A comparative phylogenomic analysis of the two *Ca*. Endonucleobacter species and 19 publicly available genomes of close relatives placed both *Ca*. Endonucleobacter species in a monophyletic clade within the family Endozoicomonadaceae (class Gammaproteobacteria), with the genus *Endozoicomonas* as their closest relatives (**Fig. 1b; Supplementary Information Table 1**). *Ca*. Endonucleobacter genomes were smaller, had reduced GC contents and considerably less amino acid synthesis pathways than related *Endozoicomonas* species and members of the non-symbiotic family Hahellaceae (**Fig. 1b**).

### *Ca*. Endonucleobacter gains its nutrition from host sugars, lipids and amino acids

How does *Ca*. Endonucleobacter gain energy and nutrition within the nucleus for its massive replication from one to more than 80,000 cells? Our metabolic reconstruction of the genomes of the two *Ca*. Endonucleobacter species, as well as the transcriptomes and proteomes of *Ca*. E. childressi revealed that nuclear DNA, RNA, and histones are unlikely to be their main source of nutrition (see next section). Instead, these intranuclear parasites likely import and consume sugars, lipids and amino acids from their host (**Fig. 2**; **Supplementary Information Tables 3, 4**). The eukaryotic nuclear pore complexes allow the passage of small molecules (≤ 30-60 kDa) between the nucleus and the cytoplasm (Knockenhauer & Schwartz, 2016; Mohr et al., 2009), providing *Ca*. Endonucleobacter with access to not only nuclear but also many cytoplasmic molecules.

**Fig. 2.**
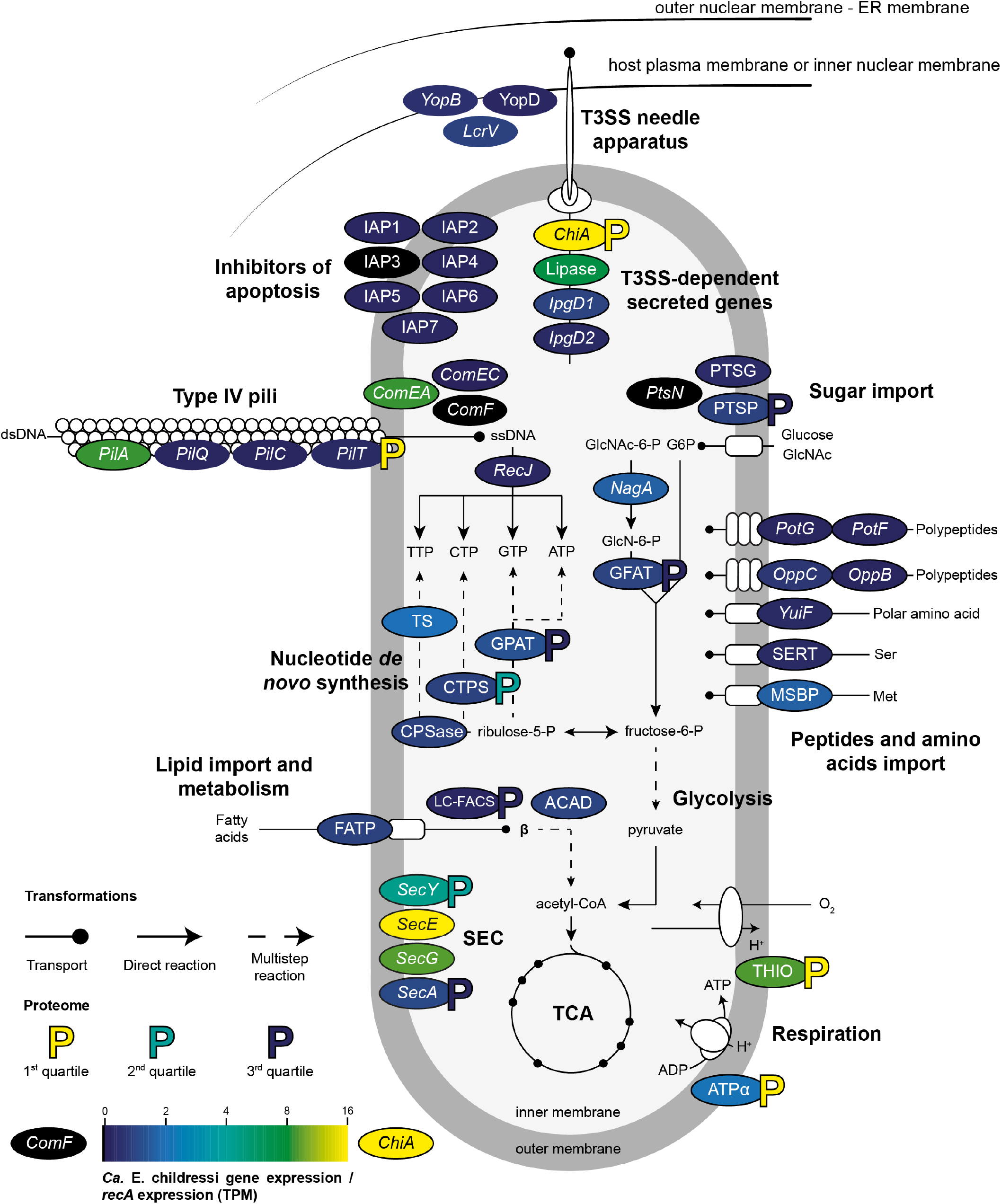
*Ca*. E. childressi is a chemoorganoheterotroph that gains it nutrition from lipids, sugars and amino acids from its host. Physiological reconstruction based on RAST annotation and Pathway Tools metabolic modelling. *Ca*. E. childressi gene expression is plotted as TPMs normalized to *RecA* TPMs. The *Ca*. E. childressi proteome is shown as colored “P symbols”, with yellow showing high abundance (first quartile), turquoise medium abundance (second quartile) and blue low abundance (third quartile). **ACAD**, acyl-CoA dehydrogenase. **ATPα**, ATP synthase alpha chain. ***ChiA***, chitinase. ***ComEA***, Late competence protein ComEA DNA receptor. ***ComEC***, DNA internalization-related competence protein ComEC. ***ComF***, competence protein F homolog. **CPSase**, carbamoyl-phosphate synthase large chain. **CTPS**, Cytidine triphosphate synthase. **FATP**, long-chain fatty acid transport protein. **GFAT**, glucosamine-fructose-6-phosphate aminotransferase. **GPAT**, amidophosphoribosyltransferase. **IAP1-7**, inhibitors of apoptosis. ***IpgD*1-2**, Shigella-like inositol phosphate phosphatases. **LC-FACS**, long-chain-fatty-acid-CoA ligase. ***LcrV***, T3SS translocon protein *LcrV*. **Lipase**, lipase. **MSBP**, methionine ABC transporter substrate-binding protein. ***NagA***, N-acetylglucosamine-6-phosphate deacetylase. ***OppB***, oligopeptide transport system permease protein *OppB*. ***OppC***, oligopeptide transport system permease protein *OppC*. ***PilA***, type IV pilin PilA. ***PilC***, Type IV fimbrial assembly protein PilC. ***PilQ***, Type IV pilus biogenesis protein PilQ. ***PilT***, Twitching motility protein PilT. ***PotF***, putrescine ABC transporter putrescine-binding protein PotF. ***PotG***, putrescine transport ATP-binding protein *PotG*. **PTSG**, glucose-specific component of PTS system. ***PtsN***, nitrogen-regulatory protein of PTS system *PtsN*. **PTSP**, phosphoenolpyruvate-protein phosphotransferase of PTS system. ***RecJ***, single-stranded-DNA-specific exonuclease *RecJ*. ***SecA***, Protein export cytoplasm protein SecA ATPase RNA helicase. ***SecE***, Preprotein translocase subunit SecE. ***SecG***, Preprotein translocase subunit SecG. ***SecY***, Preprotein translocase secY subunit. **SERT**, serine transporter. **THIO**, thioredoxin. **TS**, thymidylate synthase. ***YuiF***, histidine permease *YuiF*. ***YopB***, T3SS translocon protein *YopB*. ***YopD***, T3SS translocon protein *YopD*. Not all genes involved in *Ca*. E. childressi metabolism and pathogenesis are shown for space reasons, however, they are listed in **Supplementary tables 10 and 11**.

*Ca*. E. childressi and *Ca*. E. puteoserpentis are predicted to share highly similar metabolic pathways. They are both chemoorganoheterotrophs that encoded genes involved in glycolysis, the pentose phosphate pathway, TCA cycle and aerobic respiration with oxygen as the terminal electron acceptor (**Fig. 2**; **Supplementary Information Tables 3, 4**). Both parasites encoded lipid and sugar importers, substrates that fuel the TCA-cycle based core metabolism, and these were expressed in *Ca*. E. childressi. The two parasites lacked synthesis pathways for amino acids (10 and 11 AAs respectively) (**Fig. 1b; Supplementary Information Table 5**), but encoded importers for polypeptides such as putrescine and importers for amino acids such as the generic importer *YuiF*, which were expressed in *Ca*. E. childressi (**Fig. 2; Supplementary Information Tables 3, 4**).

Our dual RNA-seq analyses of laser microdissected early-, mid- and late infection stages of *Ca*. E. childressi revealed that it expressed nutrient importers for sugars (PTS), lipids (FATP) and amino acids (*YuiF*) throughout its infection cycle, with a particular upregulation during the early and mid infection stages (**Fig. 2, 3; Supplementary Information Table 3, 6**). Concomitantly, the host expressed genes for the import of sugars, amino acids and the synthesis of lipid droplets in the early and mid infection stages (**Fig. 3; Supplementary Information Table 7**). In the late infection stage, expression levels of genes involved in nutrient import by the parasite, as well as on the host side sugar importers and lipid droplet synthesis were lower than in the preceding stages (**Fig. 3; Supplementary Information Table 6, 7**). This could be because the host cell is no longer able to maintain its metabolism due to nutrient depletion, and/ or *Ca*. Endonucleobacter no longer grows considerably immediately before its release when the host cell bursts.

**Fig. 3.**
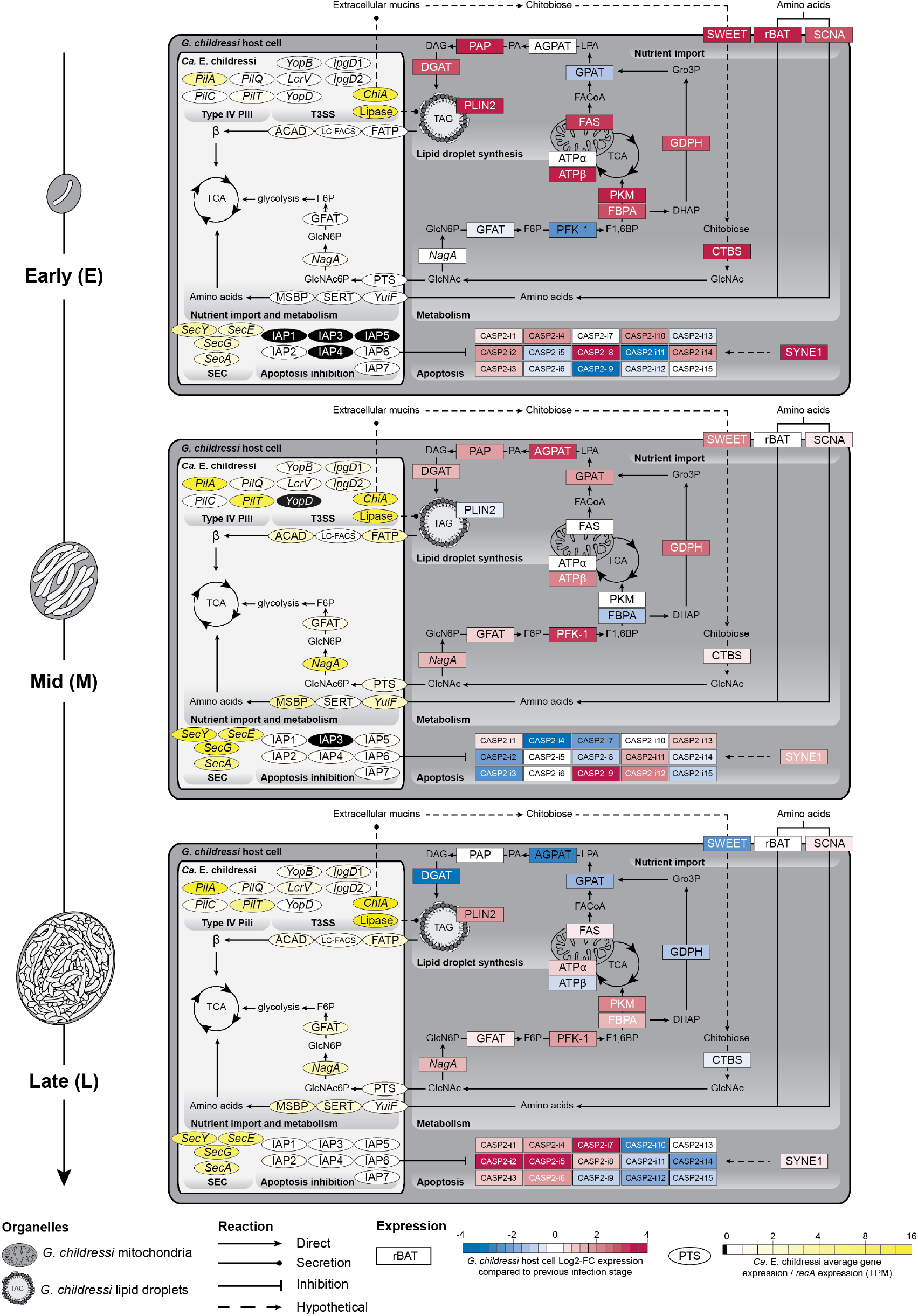
*G. childressi* gill cells remained transcriptionally and metabolically active throughout the infection cycle. In all three infection stages, inhibitors of apoptosis (IAPs) were upregulated by the parasite *Ca*. E. childressi, while the host upregulated caspases, proteins involved in initiating apoptosis that are inhibited by IAPs. Transcriptomic profiling of metabolic and apoptotic interactions between *Ca*. E. childressi (light grey) and *G. childressi* (dark grey) in early (top panel) mid, (middle panel) and late infection stages (bottom panel). *Ca*. E. childressi gene expression is plotted as average (n = 3) TPMs normalized to *RecA* TPMs. *G. childressi* gene expression is plotted as fold-changes to the previous infection stage. Fold changes of *G. childressi* gene expression at the first, early stage of infection were compared to non-infected *G. childressi* cells. ***Ca*. E. childressi genes: ACAD**, acyl-CoA dehydrogenase. ***ChiA***, chitinase. **GFAT**, glucosamine-fructose-6-phosphate aminotransferase. **IAP1-7**, inhibitors of apoptosis. ***IpgD1-2***, *Shigella*-like Inositol phosphate phosphatases. **FATP**, long-chain fatty acid transport protein. **LC-FACS**, long-chain-fatty-acid-CoA ligase. **Lipase**, probable lipase. ***LcrV***, T3SS translocon protein *LcrV*. **MSBP**, methionine ABC transporter substrate-binding protein. ***NagA***, N-acetylglucosamine-6-phosphate deacetylase. ***PilA***, type IV pilin PilA. ***PilC***, Type IV fimbrial assembly protein PilC. ***PilQ***, Type IV pilus biogenesis protein PilQ. ***PilT***, Twitching motility protein PilT. **PTS**, phosphoenolpyruvate-protein phosphotransferase of PTS system. ***SecA***, Protein export cytoplasm protein SecA ATPase RNA helicase. ***SecE***, Preprotein translocase subunit SecE. ***SecG***, Preprotein translocase subunit SecG. ***SecY***, Preprotein translocase secY subunit. **SERT**, serine transporter. ***YopB***, T3SS translocon protein *YopB*. ***YopD***, T3SS translocon protein *YopD*. ***YuiF***, histidine permease *YuiF*. ***G. childressi* genes: AGPAT**, 1-acyl-sn-glycerol-3-phosphate acyltransferase alpha. **ATPα**, ATP synthase alpha chain. **ATPβ**, ATP synthase beta chain. **CASP2-i1-15**, caspase-2 isoforms 1-15. **CTBS**, chitobiase. **DGAT**, diacylglycerol O-acyltransferase. **FAS**, fatty acid synthase. **FBPA**, fructose-bisphosphate aldolase. **GFAT**, glucosamine-fructose-6-phosphate aminotransferase. **GPAT**, glycerol-3-phosphate O-acyltransferase. **GPDH**, glycerol-3-phosphate dehydrogenase. ***NagA***, N-acetylglucosamine-6-phosphate deacetylase. **PAP**, phosphatidate phosphatase. **PFK-1**, phosphofructokinase-1. **PLIN2**, perilipin-2. **PKM**, pyruvate kinase PKM. **rBAT**, neutral and basic amino acid transport protein rBAT. **SCNA**, sodium-coupled neutral amino acid transporter. **SWEET**, SWEET sugar transporter 1. **SYNE1**, Nesprin-1. Not all genes involved in glycolysis, the TCA cycle and β-oxidation of fatty acids (β) are shown for space reasons, but are listed in **Supplementary tables 12 & 13**.

The two *Ca*. Endonucleobacter species encoded a chitinase, a trait common to many Endozoicomonadaceae (Jensen et al., 2021; Keller-Costa et al., 2022). The chitinase was highly expressed in *Ca*. E. childressi, in both the bulk transcriptomes and proteomes, as well as in the laser microdissected transcriptomes of all three infection stages (**Fig. 2, 3**; **Supplementary Information Tables 3, 4, 6**). The chitinases of both *Ca*. Endonucleobacter species encoded a signal peptide for T3SS dependent secretion, and were phylogenetically related to the ChiA-2 chitinase of *Vibrio cholerae* (**Extended Data Fig. 2**). In *Vibrio cholerae*, ChiA-2 enables it to use mucin as a source of nutrition by deglycosylating mucin and releasing sugars like N-acetylglucosamine (GlcNAc) and its oligomers (Mondal et al., 2014). If ChiA-2 functions similarly in *Ca*. E. childressi, extracellular mucins of the mussel, produced by the secretory cells of the gill (**Supplementary figure 1**), would provide a rich source of nutrition, and their uptake could occur through the upregulated importers in both the host and the parasite for sugars. However, it is not clear if the chitinase could be secreted by the T3SS through the nuclear envelope into the host cytoplasm, and how it would then be exported through the host epithelial membrane to the extracellular mucin.

### *Ca*. Endonucleobacter does not appear to gain nutrition from DNA, RNA or histones

*Ca*. Endonucleobacter is unlikely to use DNA, RNA and histones as its main source of nutrition based on the following evidence. Both *Ca*. Endonucleobacter species lacked genes needed for external secretion of DNAses, RNAses or proteases (**Supplementary Information Table 1**). When using DNA for nutrition, *V. cholerae* or *Escherichia coli* secrete DNAses to the extracellular medium or the periplasm, where the DNA is digested extracellularly and oligonucleotides and monomers imported by the cell (Huang et al., 2022; Seper et al., 2011). While *Ca*. E. childressi expressed DNAses such as exodeoxyribonucleases I, III, V, VII and RecJ, none of them were predicted to have secretion signal peptides, indicating that these DNAses are used for housekeeping tasks like DNA replication and repair and recycling. Nucleotide importers such as NupC, which is used by *V. cholerae* and *E. coli* (Huang et al., 2022; Seper et al., 2011), were absent in both *Ca*. Endonucleobacter species. Furthermore, the two parasites had all the genes needed for synthesizing their own nucleotides, and they were expressed in *Ca*. E. childressi (**Fig. 2**; **Supplementary Information Tables 3, 4, 8**).

Both *Ca*. Endonucleobacter species encoded competence factors for DNA import such as *comEA, comEC* and *comF*, but only *comEA* was highly expressed in *Ca*. E. childressi, *comEC* was expressed at low levels and *ComF* not at all (**Fig. 2**; **Supplementary Information Tables 3, 4**). While Type IV pili (T4P) can play a role in DNA uptake (Ellison et al., 2022), and most T4P genes were expressed in *Ca*. E. childressi (**Fig. 2**; **Supplementary Information Tables 3, 4, 9**), T4P are also known to play roles in facilitating adherence to host cells, surface movement (twitching motility), phage adsorption and biofilm formation (Craig et al., 2019; Evans et al., 2007; Matthey & Blokesch, 2016; Medina et al., 2008).

In summary, while we cannot exclude that T4P and competence genes could be involved in DNA uptake in *Ca*. Endonucleobacter, the lack of expression of genes involved in importing nucleotides and secreting nucleases, together with the high expression of genes involved in the digestion of sugars, lipids and amino acids, indicates that DNA is not the main source of nutrition for *Ca*. E. childressi (**Fig. 2; Supplementary Information Tables 3, 4**).

On the host side, our analyses of different infection stages provided further evidence that *Ca*. E. childressi does not appear to consume considerable amounts of host DNA, RNA or histones (**Supplementary Information Table 7**). Our dual RNA-seq analyses of laser microdissected infection stages revealed no evidence for downregulation of host transcription, as expected if nuclear DNA and RNA were consumed (Schulz and Horn 2015). The infected host cell remained transcriptionally and metabolically active throughout the infection cycle, as host genes involved in glycolysis and oxidative phosphorylation were expressed, even in late-stage nuclei (**Fig. 3; Supplementary Information Table 7**). Moreover, light and electron microscopy analyses revealed that the mussel host cells remained morphologically asymptomatic, apart from the swollen nucleus, with intact membranes and organelles (**Extended Data Fig. 3, 4**).

### *Ca*. Endonucleobacter and its mussel host engage in an apoptotic arms race

One highly unusual feature of *Ca*. Endonucleobacter is that its genome encodes inhibitors of apoptosis (IAPs), with *Ca*. E. childressi and *Ca*. E. puteoserpentis encoding seven and 13 IAPs respectively (**Supplementary Information Table 1**; **Supplementary Note 2**). IAPs are an evolutionarily conserved group of proteins known from animals and viruses that have not been previously described in bacteria. In animals, IAPs inhibit a process of programmed cell death called apoptosis, mainly by binding caspases, proteases that play a central role in executing apoptosis (reviewed in Ramirez & Salvesen, 2018; Salvesen & Duckett, 2002). Members of the IAP protein family contain one to three baculoviral IAP repeat (BIR) motifs that allow them to sequester caspases (Salvesen & Duckett, 2002). IAPs are sometimes referred to as BIRPs (BIR-containing proteins), as not all proteins of the IAP family are bona fide inhibitors of apoptosis (Uren et al., 1998). Only BIRPs that have a RING domain, which can ubiquinate caspases to target them for proteolysis via the proteasome, are considered bona fide apoptosis inhibitors (Ni et al., 2005; Vaux & Silke, 2005). In our analysis, we therefore only refer to proteins as IAPs if they had both a BIR and a RING domain (**Extended Data Fig. 5**).

To understand the role IAPs play in the biology of *Ca*. Endonucleobacter, we studied the genes expressed by both the parasite and its host during the infection cycle (**Fig. 2, 3; Supplementary Information Table 3, 4, 6, 7**). All seven IAPs encoded by *Ca*. E. childressi had signal peptides for the Sec secretion pathway, and the genes *SecA, SecY, SecE* and *SecG* were expressed in all infection stages (**Fig. 2, 3; Supplementary Information Table 3, 4, 6**). *Ca*. E. childressi first expressed three IAPs in early infection stages, then six in mid stages, and finally all seven IAPs in late stages of infection (**Fig. 3; Supplementary Information Table 6**). Concomitantly, the host expressed as many as 16 different capase-2 isoforms throughout the infection cycle (**Fig. 3; Supplementary Information Table 7**).

A wide range of stimuli can trigger apoptosis, such as metabolic stress, DNA damage and ER stress. One of the caspases that initiates the apoptotic cascade is caspase-2 (reviewed in Kopeina & Zhivotovsky, 2021). IAPs are well known for their ability to bind to and block caspases (Salvesen & Duckett, 2002). Although not well studied in marine invertebrates, in the oyster *Crassostrea gigas*, IAP-2 strongly binds to and blocks caspase-2, suggesting that IAPs play an important role in the inhibition of caspase-2-mediated apoptosis in bivalves (Qu et al., 2015). The concomitant upregulation during the infection cycle of host caspase-2 isoforms by *G. childressi* and bacterial IAPs by *Ca*. E. childressi suggest that the host initiates apoptosis in response to the infection, dramatic swelling of its nucleus and high-jacking of its metabolism (**Supplementary Note 3, 4**), which *Ca*. E. childressi encounters by upregulating IAPs. Thus, both the host and the intranuclear parasite engage in a physiological arms race for control of apoptosis, with seven different IAPs of *Ca*. E. childressi preventing an arsenal of *G. childressi* caspase isoforms from inducing apoptosis long enough for the parasite to acquire the energy and nutrients it needs to replicate to such high numbers before the death of its host cell.

### *Ca*. Endonucleobacter acquired inhibitors of apoptosis from its mussel host

Despite the fact that IAPs have not been previously reported from bacterial genomes, our analyses revealed that in addition to *Ca*. Endonucleobacter, four *Endozoicomonas* species, isolated from other marine invertebrates, encoded bona fide IAPs (**Supplementary Information Table 1**). Comparative phylogenetic analyses of *Ca*. Endonucleobacter and other Endozoicomonadaceae IAPs with publicly available animal and viral IAPs revealed that bacterial IAPs were not monophyletic, but rather fell into seven clades that were interspersed with IAPs of marine invertebrates (**Fig. 4**). IAPs in all three *Ca*. Endonucleobacter clades were most closely related to those of their hosts, as well as other molluscs (**Fig. 4**). Similarly, another group of bacterial IAPs, from *Endozoicomonas* isolated from ascidians and called *Endozoicomonas ascidiicola*, were most closely related to ascidian IAPs (**Fig. 4**). Two viral IAPs from the ostreid herpes virus OsHV-1 were also interspersed across the tree, and more closely related to bacterial and invertebrate IAPs than to each other. OsHV-1, first found in oysters, infects a wide range of molluscs (Arzul et al., 2001a; Arzul et al., 2001b; Hine et al., 1998; Hine & Thorne, 1997; Prado-Alvarez et al., 2021), including *G. childressi*, based on the recovery of 17% of the OsHV-1 genome in one of the *G. childressi* specimens sequenced in this study (**Supplementary Note 5**).

**Fig. 4.**
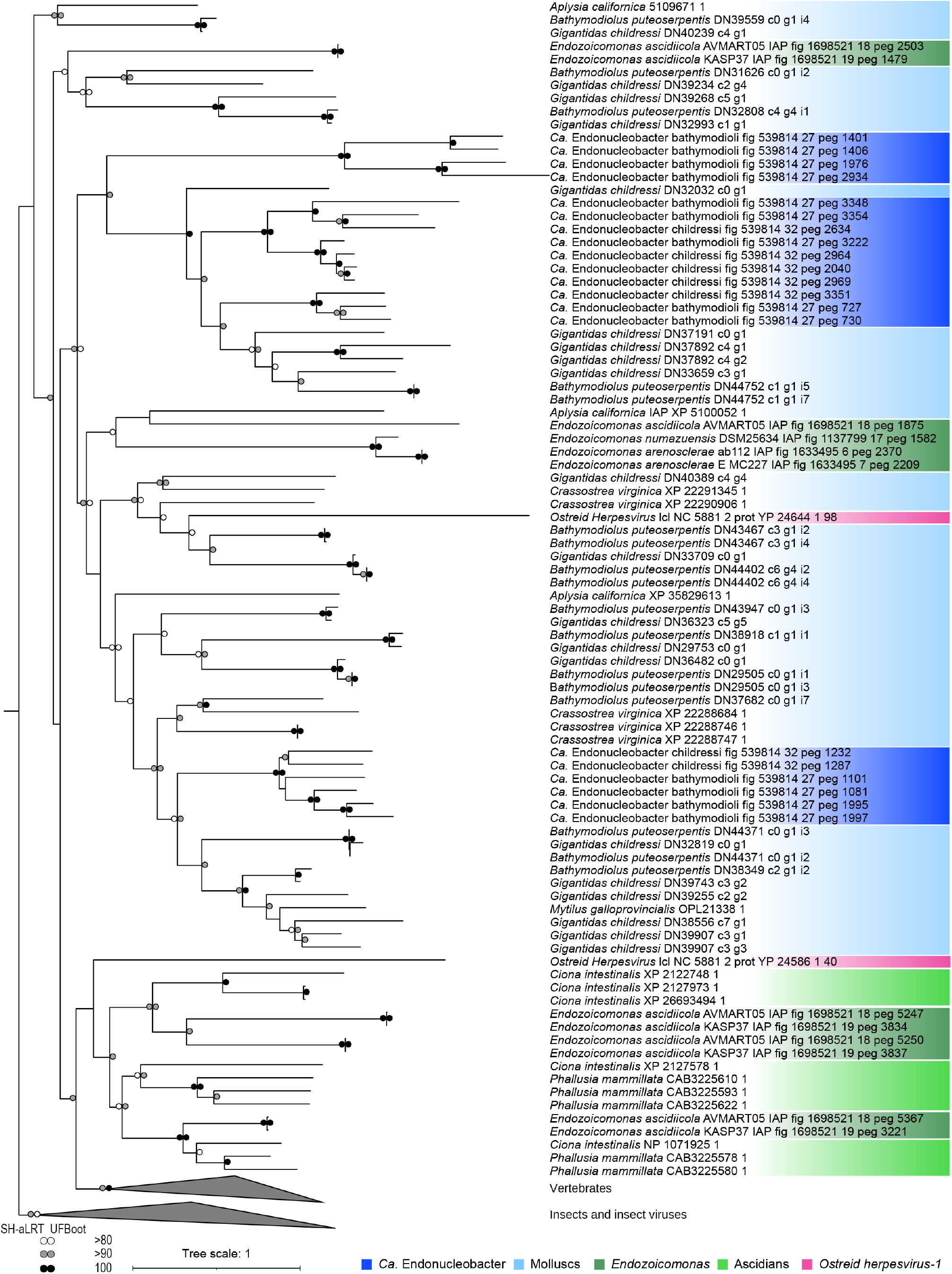
IAPs encoded by *Ca*. Endonucleobacter are interspersed with those of their mussel host and ostreid herpes viruses. Protein-based phylogeny of 128 IAPs from *Ca*. Endonucleobacter, *Endozoicomonas*, metazoans and viruses (bootstrap: 1000). Bootstrap support values ≥ 80 are shown. Scale bar indicates substitutions per site. Colors indicate taxonomic groups. Insects and their virus IAPs were included to root t he tree. Triangle length indicates number of sequences contained within collapsed clades.

The interspersed phylogeny of Endozoicomonadaceae IAPs with those of marine invertebrate and viral IAPs suggests horizontal gene transfer (HGT) of these genes between animals, bacteria and viruses. Given that apoptosis is specific to animals, and IAPs are only known from animals and a few viruses that infect insects and molluscs (Davison et al., 2005; Huang et al., 2000; Hughes, 2002; Nagamine, 2022), IAPs in bacteria are likely not ancestral, but were rather acquired through HGT from animals or viral vectors. HGT from animals to bacteria has only rarely been observed, although it is common between bacteria, and from bacteria to eukaryotes, particularly in protists (Keeling & Palmer, 2008). One explanation for why eukaryote to prokaryote HGT appears to be rare is that sequestration of eukaryotic DNA within the nucleus makes it harder to access. This would not be the case for *Ca*. Endonucleobacter, whose intranuclear lifestyle places it in direct contact with its eukaryotic host’s DNA.

As to how *Ca*. Endonucleobacter acquired its host’s IAPs, three mutually non-exclusive explanations are plausible: (*i*) *Ca*. Endonucleobacter acquired the IAPs via its competence genes or T4P for taking up DNA (**Fig. 2**). (*ii*) Horizontal gene transfer of IAPs could have also been facilitated by the numerous mobile elements in the genomes of *Ca*. Endonucleobacter (**Supplementary Information Table 1**). (*iii*) Viruses that infect bacteria living inside eukaryotic cells can act as vectors, as shown in some insects that have acquired bacterial toxins through HGT from phages that infect their symbiotic bacteria (Verster et al., 2021) (**Supplementary Note 5**). Our sequencing analyses revealed that *Ca*. E. childressi encoded 2 CRISPR-Cas spacers with similarity to the genomes of Herpesvirales, viruses so far only known to infect eukaryotes, including two CRISPR-Cas spacers that mapped to OsHv-1 with 65.6% (intergenic region) and 62.5% (BIR motif) (**Supplementary Fig. 2**; **Supplementary Information Table 14**). This indicates that *Ca*. E. childressi recently encountered herpes viruses, as bacteria insert CRISPR-Cas spacers specific to the virus infecting them as part of their immune response. Moreover, our TEM analyses revealed viral structures inside *Ca*. E. childressi (**Extended Data Fig. 6**).

## Conclusion

*Ca*. Endonucleobacter and *Endozoicomonas* IAPs are more closely related to IAPs from their animals hosts than to each other, and fall into clades separated by IAPs from other animals, indicating that eukaryote to bacteria HGT of IAPs occurred more than once in the Endozoicomonadaceae (**Fig. 4**). IAPs have ankyrin repeats (ANKs) in their RING domain, a protein motif that is widespread and common in eukaryotes and proposed to have been acquired by bacteria through HGT (Bork, 1993). Like *Ca*. Endonucleobacter, *Endozoicomonas* genomes are enriched in ANKs and mobile elements, suggesting that these genomic features are a common trait of the family Endozoicomonadaceae (**Supplementary Information Table 1)** (Alex & Antunes, 2019; Neave et al., 2017; Pogoreutz et al., 2022, Ide et al., 2022). Mobile elements are known to promote HGT in microorganisms (Haudiquet et al., 2022), and might have facilitated repeated acquisition of IAPs and other animal host genes, enriching Endozoicomonadaceae genomes with these eukaryotic-like domains. Given that eukaryotic-like domains are enriched in bacteria associated with eukaryotes, together with studies showing that these proteins modulate interactions between bacteria and their eukaryotic hosts, it is possible that genomic enrichment in ANKs and other eukaryotic-like proteins enabled the enormous versatility of Endozoicomonadaceae in associating with a wide range of animals from marine environments around the world (Ding et al., 2016; Gomez-Valero et al., 2011; Maire et al., 2023; Nguyen et al., 2014) (reviewed in Neave et al., 2016).

The presence of IAPs in an intranuclear bacterial parasite poses the chicken or the egg question. Did the intranuclear lifestyle of *Ca*. Endonucleobacter allow the acquisition of IAPs from its host, or did the acquisition of IAPs allow *Ca*. Endonucleobacter to make a living in the nucleus? Acquisition of host IAPs by *Ca*. Endonucleobacter would have required at least two steps; i) intimate contact with host’s DNA or mRNA or that of a virus of the host, and ii) a certain frequency of this contact. Both requirements would suggest an intracellular lifestyle with the ability to invade the nucleus at least occasionally. *Rickettsia* that infect insects have such a lifestyle, as they are generally intracellular but occasionally invade their host’s nuclei (Burgdorfer et al., 1968; Burgdorfer & Brinton, 1970; Kuechler et al., 2013; Perotti et al., 2006; Simser et al., 2002; Whitworth et al., 2003). The presence of IAPs in *Endozoicomonas*, some of which cluster with the IAPs from their ascidian host, raises the question if these *Endozoicomonas* could also occasionally invade their host’s nuclei. To date, most Endozoicomonadaceae associations have not been analyzed with imaging methods that would resolve this question.

Not all intranuclear bacteria use IAPs to avoid apoptosis. For example, the intranuclear bacteria that colonize microbial eukaryotes do not encode IAPs. However, microbial eukaryotes lack bona fide caspases, and in some of them, apoptosis is caspase independent (Olie et al., 1998; Saheb et al., 2014). Moreover, if a bacterium that lives in a unicellular host is passed on to both daughter cells, it could lead a sheltered, intranuclear lifestyle as long as the bacterium does not have major negative effects on its host’s fitness. In contrast, the intranuclear bacteria of deep-sea mussels colonize terminally differentiated cells and must therefore reproduce before their host cell dies, explaining the strong selective advantage in acquiring IAPs.

The lack of eukaryotic metabolic versatility compared to bacteria is one explanation why HGT from eukaryotes to prokaryotes is so rare (Keeling & Palmer, 2008). However, eukaryotes have evolved numerous genes and pathways for interacting with both beneficial and parasitic bacteria. Acquisition of these genes by bacteria could improve their ability to enter and reproduce in their eukaryotic hosts, as argued here for the acquisition of IAPs by *Ca*. Endonucleobacter. One of the most striking examples for the selective advantage of having eukaryotic-like proteins is *Legionella pneumophila*, which acquired a large number of these proteins as effectors for interfering with host pathways (Cazalet et al., 2004; Gomez-Valero et al., 2019). Similarly, work on sponges identified eukaryotic-like proteins in their symbionts that mediate phagocytosis (Reynolds & Thomas, 2016). These examples indicate that HGT from eukaryotes to bacteria may be more common than currently recognized, particularly in bacteria that are closely associated with eukaryotic hosts (Martyn et al., 2022). As large-scale sequencing efforts aimed at a holistic view of the genomic underpinnings of eukaryotic organisms and their associated microbiome are now becoming more common, we are in an ideal position to revisit our understanding of eukaryote-to-prokaryote HGT events.

## Material and methods

### Sample collection

*Gigantidas childressi* mussels were collected with the remote operated vehicle Hercules during the RV Meteor Nautilus NA-58 cruise to the Gulf of Mexico in May 2015. *G. childressi* mussels were collected at the Mississippi Canyon site (MC853, 28º07’ N; -089º08’ W) and the Green Canyon site (GC234, 27º45’ N; -091º13’ W) at water depths of 1,070 and 540 m, respectively. *B. puteoserpentis* mussels were collected with the remote operated vehicle MARUM-QUEST during the during the Sonne M-126 cruise to the Mid-Atlantic Ridge in April 2016. *B. puteoserpentis* mussels were collected from the Irina II venting site (Irina-II, 14º45’ N; -044º59’ W) at a water depth of 3,036 m. Upon recovery, the gills of the collected specimens were dissected on board, preserved and stored according to their final purpose. Samples for DNA sequencing, bulk RNA sequencing and polyA-RNA sequencing were preserved in RNA*later* (Thermo Fisher Scientific, MA, USA) and stored at -80°C. Samples for microscopy and laser-capture microdissection transcriptomic analyses were fixed in paraformaldehyde (PFA) 4% in 1X PBS for 8 h at 4 °C and stored in 0.5 X PBS – 50 % ethanol at -20 °C. Samples for proteomics were snap frozen with liquid nitrogen and stored at -80 °C. Samples for transmission electron microscopy were fixed in 2.5% glutaraldehyde (GA) in PHEM buffer (piperazine-N, N′-bis, 4-(2-hydroxyethyl)-1-piperazineethanesulfonic acid, ethylene glycol-bis (β-aminoethyl ether and MgCl2 (Montanaro et al. 2016)). After fixation, samples were stored in PHEM buffer. Sampling metadata of collected specimens can be found in **Supplementary information table 15**.

### Microscopy

#### Whole-mount fluorescence *in situ* hybridization

Three gill filaments of *G. childressi* (H1423/002-N9) and of *B. puteoserpentis* (499ROV/1-4) were dissected and hybridized with 16S rRNA-targeting probes in hybridization buffer. Probes used in this study are listed in **Supplementary information table 2**. The hybridization buffer contained 35% formamide, 80 mM NaCl, 400 mM Tris HCl, 0.4% blocking reagent for nucleic acids (Roche, Switzerland)), 0.08% SDS (v/v) and 0.08 dextran sulfate (w/v). We dissolved 5 ng·µl^-1^ of the Eubacterial probe EUB-I 338 (5’GCTGCCTCCCGTAGGAGT3’) (Amann et al., 1990) and 5 ng·µl^-1^ of the *Ca*. Endonucleobacter probe BNIX64 (5’GCTAGACCTGTTACCGCT3’) (Zielinski et al., 2009) in hybridization buffer. We hybridized the gill filaments in 50 µL of this hybridization solution at 46°C for 3 h. Following hybridization, the gill filaments were washed in pre-warmed 48°C washing buffer (0.07 M NaCl, 0.02 M Tris-HCl pH 7.8, 5 mM EDTA pH 8, and 0.01% SDS (v/v)) for 15 min. After washing, the gill filaments were counterstained with DAPI for 10 min at room temperature, transferred to poly-L-lysine-coated glass slides (Sigma-Aldrich, MO, USA) and mounted using the ProLong^®^ Gold antifade mounting media (Thermo Fisher Scientific, MA, USA), cured overnight at room temperature and stored at -20°C until visualization.

#### Fluorescence microscopy

Whole-filament overviews (**Extended Data Fig.1, a & d**) were visualized with the epifluorescence microscope Olympus BX53 (Olympus, Germany) with a UCPlanFL 20X/0.70 air transmission lens and an Orca Flash 4.0 camera (Hamamatsu, Japan) using the Olympus cellSens Dimension software v. 1.18 (Olympus, Germany). Detailed images (**Fig.1, a**), (**Extended Data Fig.1, b-c & e-f**), (**Supplementary Fig. 1, b & c**) and (**Extended Data Fig. 4, a-c**) were recorded with a Zeiss LSM 780 equipped with an Airyscan detector and two different objectives, a plan-APROCHROMAT 63X/1.4 oil immersion objective and a plan-APROCHROMAT 100X/1.46 DIC M27 Elyra oil immersion objective. Images were obtained and post-processed using ZEN software (black edition, 64bits, version: 14.0.1.201) (Carl Zeiss Microscopy GmbH, Germany). Images were brightness and levels-adjusted using the software Adobe Photoshop (version: 12.0) (Adobe Systems, CA, USA).

#### N-acetylglucosamine-based saccharides localization

To identify saccharides potentially targeted by *Ca*. E. childressi chitinase, we localized N-acetylglucosamine residues in *G. childressi* gill tissues (**Supplementary Fig. 1**). The formalin-fixed paraffin-embedded gills of the *G. childressi* specimen H1423/001-N5-002 were sectioned at 10 µm thickness using a microtome. Sections were mounted on poly-L-lysine-coated glass slides, and left to dry at RT for 4 h. Sections were baked at 60°C for 1 h to improve tissue adherence and dewaxed with three 10 min steps of Roti^®^-Histol (Carl-Roth, Germany) followed by a decreasing ethanol series of 96, 80, 70 and 50% (v/v) for 10 min each. Then, tissue sections were washed in milliQ water for 10 min, and dried at 37 °C for 30 min. Hybridization was performed as described in the section above, with the BNIX64 probe. After hybridization, sections were washed 1 min in milliQ water, dipped into ethanol 96% (v/v) and dried at 37°C for 30 min. The Fungi-Fluor^®^ Kit (Polysciences, PA, USA) was applied following the manufacturers protocol. All sections were mounted using the ProLong^®^ Gold antifade mounting media (Thermo Fisher Scientific, MA, USA), cured overnight at room temperature and stored at -20°C until visualization.

#### *G. childressi* host cell 18S rRNA localization

Gill sections from the *G. childressi* specimen H1423/001-N5-002 were hybridized with the probe BNIX64, and the eukaryotic EUK-1195 probe (5’GGGCATCACAGACCTG’3) (Giovannoni et al., 1988) (**Extended Data Fig. 4, a-c)**. The 18S rRNA fluorescence intensity of *G. childressi* cells at different the different stages of *Ca*. E. childressi infection (Non-infected, NI; early-stage of infection, E; mid-stage of infection, M; late-stage of infection, L) was measured in ten areas of identical area, using Fiji (Schindelin et al., 2012) and the average fluorescence intensity plotted in **Extended Data Fig. 4, d**.

#### Transmission electron microscopy

Samples in (**Fig.1, a**) and (**Extended Data Fig. 3 & 6**) were post fixed with 1% (v/v) osmium tetroxide (OsO4) for 2 h at 4 °C, washed three times with PHEM and dehydrated with an ethanol series (30%, 50%, 70%, 80%, 90% and 100% (v/v)) at -10 °C, each step lasting 10 min. The samples were then transferred into 50:50 ethanol and acetone, followed by 100% acetone and infiltrated with low-viscosity resin (Agar Scientific, UK) using centrifugation embedding (McDonald, 2014). Samples were centrifuged for 30s in resin: acetone mixtures of 25%, 50%, 75% and twice in 100%. Finally they were transferred into fresh resin in embedding molds and polymerized at 60–65 °C for 48 h. Ultra-thin (70 nm) sections were cut on a microtome (Ultracut UC7 Leica Microsystem, Austria), mounted on formvar-coated slot grids (Agar Scientific, United Kingdom) and contrasted with 0.5% aqueous uranyl acetate (Science Services, Germany) for 20 min and with 2% Reynold’s lead citrate for 6 min. Sections were imaged at 20–30 kV with a Quanta FEG 250 scanning electron microscope (FEI Company, USA) equipped with a STEM detector using the xT microscope control software ver. 6.2.6.3123.

### Proteomics

#### Proteomics sample preparation

We dissected the ciliated edge from 20 snap frozen gill filaments of 12 *G. childressi* specimens: H1423/001-N1, H1423/001-N2, H1423/001-N5, H1423/002-N9, H1423/002-N10, H1423/003-N11, H1423/003-N15, H1423/004-N19, H1425/019-N59, H1425/019-N60, H1425/019-N61 and H1425/019-N62. We conducted a tryptic protein digestion following the filter-aided sample preparation (FASP) protocol, adapted from Wiśniewski et al. 2009 for all samples. Depending on the amount of available tissue, 100 or 150 µl of SDT-lysis buffer (4% (w/v) SDS, 100 mM Tris-HCl pH 7.6, 0.1 M DTT) were added and boiled samples at 95 °C for 10 min. To minimize sample loss, we did not do the 5 minutes centrifugation step at 21,000g as described in the original FASP protocol Wiśniewski et al. 2009. Instead, only a short spin down was conducted. Subsequently, lysate and UA solution were mixed in a 3:10 ratio (8 M urea in 0.1 M Tris/HCl pH 8.5) in a 10 kDa MWCO 500 µL centrifugal filter unit (VWR International) and the mixture was centrifuged at 14,000g for 40 min. This step was repeated multiple times until the whole lysate was loaded onto the filter unit. Next, we added 200 µL of UA solution and centrifugal filter spun again at 14,000g for 40 min. Subsequently, 100 µL of IAA solution (0.05 M iodoacetamide in UA solution) were added and samples were incubated at 22 °C for 20 min in the dark. We removed the IAA solution by centrifugation following three washing steps with 100 µL of UA solution. Subsequently, filters were washed three times with 100 µL of ABC buffer (50 mM ammonium bicarbonate). We added 1.05 μg of Pierce MS grade trypsin (Thermo Fisher Scientific, Germany) in 40 µL of ABC buffer to each filter. Filters were incubated overnight in a wet chamber at 37 °C. The next day, we eluted the peptides by centrifugation at 14,000g for 20 min followed by addition of 50 µL of 0.5 M NaCl, shaking at 600 rpm and another centrifugation step. Peptides were quantified using the Pierce MicroBCA Kit (Thermo Fisher Scientific, Germany) following the instructions of the manufacturer. For each run, 1500 ng of peptide solution were loaded onto a 5 mm, 300 µm ID C18 Acclaim® PepMap100 pre-column (Thermo Fisher Scientific, Germany) using an UltiMate™ 3000 RSLCnano Liquid Chromatograph (Thermo Fisher Scientific, Germany) and desalted on the pre-column. After desalting the peptides, the pre-column was switched in line with a 75µm x 75 cm analytical EASY-Spray column packed with PepMap RSLC C18, 2 µm material (Thermo Fisher Scientific, Germany), which was heated to 55 °C. The analytical column was connected via an Easy-Spray source to a Q Exactive HF-X Hybrid Quadrupole-Orbitrap mass spectrometer (Thermo Fisher Scientific, Germany). Peptides were separated on the analytical column using a 460 min gradient as described by Kleiner et al. 2017 with the modification that the gradient went from 98% buffer A (0.1% formic acid) to 31% buffer B (0.1% formic acid, 80% acetonitrile) in 200 min, then from 31% to 50% buffer B in 40 min and ending with 20 min at 99% buffer B. Mass spectra were acquired in the Orbitrap as described by Hinzke et al. 2019 with some modifications. In brief, eluting peptides were ionized via electrospray ionization (ESI) and analyzed in Q Exactive HF-X. Full scans were acquired in the Orbitrap at 60,000 resolution. The 15 most abundant precursor ions were selected in a data dependent manner, isolated with the quadrupole with a 1.2 m/z isolation window size, fragmented in the HCD with 25 NCE, and measured in the Orbitrap at 7,500 resolution. The mass (m/z) 445.12003 was used as lock mass as described in Olsen et al. 2005 with the modification that lock mass was detected in the full scan rather than by separate selected ion monitoring scan injection. Lock mass use was set to ‘best’. Singly charged ions were excluded from MS/MS analysis. Dynamic exclusion was set to 30 s. On average, 258,842 MS/MS spectra were acquired per sample with the 460 min gradient.

#### Proteomics data processing

A database containing protein sequences from *Ca*. E. childressi genome was used. The cRAP protein sequence database (http://www.thegpm.org/crap/) containing protein sequences of common laboratory contaminants was appended to the database. The final database contained 38,281 protein sequences. Searches of the MS/ MS spectra against this database were performed with the Sequest HT node in Proteome Discoverer version 2.2.0.388 (Thermo Fisher Scientific, Germany) as described previously in Gruber-Vodicka et al. 2019. Only proteins identified with medium or high confidence were retained resulting in an overall false discovery rate of <5%. For protein quantification, normalized spectral abundance factors (NSAFs, Zybailov et al. 2006, Nucleic Acids Research, Volume 44, Issue D1) were calculated per species and multiplied by 100, to give the relative protein abundance in %. *Ca*. E. childressi biologically meaningful factors from the global bulk proteomic analysis from the *G. childressi* specimen H1423/002-N9 (highlighted in green in **Supplementary information table 11)** were used to complement the metabolic reconstruction of *Ca*. E. childressi (**Figure 2; Supplementary information table 4**).

### DNA and RNA extraction

#### DNA extraction and infection screening

We PCR screened 14 *G. childressi* and 5 *B. puteoserpentis* RNA*later*-preserved gill samples for *Ca*. Endonucleobacter infection (**Supplementary information table 15**). DNA was extracted using the DNeasy Blood and Tissue Kit (QIAGEN, Germany) following manufacturer’
ss protocol and used as template in PCR reactions. The *Ca*. Endonucleobacter 16S rRNA gene was PCR-amplified using the following conditions: Initial denaturation for 3 min at 95 °C, followed by 30 cycles at 95 °C for 30 s, 55 °C for 30 s and 72 °C for 2 min, followed by a final elongation step at 72 °C for 10 min. The *Ca*. Endonucleobacter 16S rRNA gene was amplified using the forward primer BNIX64 (AGCGGTAACAGGTCTAGC) (Zielinski et al., 2009) and the reverse primer GM4 (TACCTTGTTACGACTT) (Muyzer et al., 1995). We used Taq DNA Polymerase (5 PRIME, Hamburg, Germany) in all PCR reactions. We loaded PCR products in agarose gels (2%) and stained with ethidium bromide for 30 min, considering band thickness as indicator of infection degree.

#### Metagenomic library preparation and sequencing

We sequenced the DNA of one specimen each of *G. childressi* (H1423/002-N9) and *B. puteoserpentis* (499ROV/1-4) specimens using short-read (Illumina HiSeq 3000) and long-read (PacBio) sequencing technologies. Sequencing was done at the Max Planck Genome Center Cologne, Germany (https://mpgc.mpipz.mpg.de/home/). For short-read sequencing, 50 ng of genomic DNA were fragmented via sonication (Covaris S2, Covaris), followed by library preparation with NEBNext Ultra DNA v2 Library Prep Kit for Illumina (New England Biolabs). Library preparation included 7 cycles of PCR amplification. Quality and quantity were assessed at all steps via capillary electrophoresis (TapeStation, Agilent Technologies) and fluorometry (Qubit, ThermoFisher Scientific). The library was immobilized and processed onto a flow cell with cBot (Illumina) and subsequently sequenced on HiSeq3000 system (Illumina) with 2 x 150 bp paired end reads. 333 million paired-end reads were generated.

Long-read sequencing was done according to the manual “Procedure-Checklist-20-kb-Template-Preparation-Using-BluePippin-Size-Selection” of Pacific Biosciences without initial DNA fragmentation and without a final size-selection. Instead, library was finally purified twice with PB AMPure beads. Sequencing was performed on a Sequel device with Sequel Binding Kit 3.0 and Sequel Sequencing Kit 3.0 for 20 h (Pacific Biosciences). A total of three sequencing PacBio cells were generated.

#### *G. childressi de novo* transcriptome: RNA extraction, library preparation and sequencing

To study the host cell expression along the infectious cycle, we assembled a *G. childressi de novo* transcriptome. We dissected the ciliated edge from 20 RNAlater-preserved gill filaments of the *G. childressi* H1423/002/N6. RNA was extracted and prepared as described in the section “Bulk transcriptomics: RNA extraction and sequencing” with the following modifications: 1 µg of total RNA was used for library preparation, polyA enrichment was done with the NEBNext poly(A) mRNA Magnetic Isolation Module (New England Biolabs), library preparation was done with the NEBNext Ultra II Directional RNA Library Prep Kit for Illumina (New England Biolabs), library preparation included 11 cycles of PCR amplification. A total of 99 million paired-end reads were generated.

#### Bulk transcriptomics: RNA extraction and sequencing

We dissected the ciliated edge from nine RNA*later*-preserved gill filaments of the *G. childressi* specimen H1423/002-N9 and extracted total RNA using the RNeasy Mini Kit (QIAGEN, Germany) following manufacturer’
ss protocol. RNA quantity was measured with a QUANTUS Fluorometer (Promega, Germany). Library preparation and sequencing was performed as described in the section “DNA sequencing and hybrid genome assembly” for the short-read library preparation, with the following modifications: 20 ng of total RNA was used for library preparation, library preparation was done with the NEBNext Ultra II Directional RNA Library Prep Kit for Illumina (New England Biolabs). A total of 33 million paired-end reads were generated.

### Laser capture microdissection

#### Dissection of infected nuclei of *G. childressi*

We used the formalin-fixed gills of the mussel H1423/002/N6 for laser-capture microdissection. We embedded the gill filaments in polyester wax, sectioned them at 10 µm thickness using a microtome and mounted the sections on thermoexitable POL-membranes (No. 115005191; Leica, Germany). Next, we hybridized the sections using the *Ca*. Endonucleobacter 16S rRNA probe BNIX64. Hybridization was performed as described earlier with the following modifications: The hybridization buffer did not contain formamide, only the *Ca*. Endonucleobacter 16S rRNA was hybridized, sections were not DAPI-stained and no mounting medium was used after air-drying. A Leica LMD6500 (Leica, Germany) was used to dissect the hybridized samples. Per infection stage, 100 nuclei were microdissected and pooled in a single tube prefilled with 30 µL of extraction buffer (AllPrep DNA/RNA FFPE kit; Qiagen, Germany). Additionally non-infected nuclei were retrieved to establish a baseline of host expression. For each condition, triplicates were taken, resulting in a total of 1,200 micro-dissected nuclei in 12 samples.

#### LCM transcriptomics: RNA extraction, library preparation and sequencing

We extracted RNA from the micro-dissected nuclei using the AllPrep DNA/RNA FFPE kit (Qiagen, Germany) following manufacturer protocol with the following modifications: Samples were incubated in proteinase K overnight at 37°C. Elution buffer was pre-warmed at 37°C and added to the column membrane. Elution buffer incubation time was doubled. After a first elution step, the eluent was loaded again on the membrane, incubated for 2 minutes and eluted again. RNA quantity was assessed with a QUANTUS Fluorometer (Promega, Germany). Library preparation and sequencing was done as described in the section “DNA sequencing and hybrid genome assembly” for the short-read library preparation, with the following modifications: total RNA was amplified following the protocol of capture and amplification by tailing and twitching described in Turchinovich et al. 2014. Library preparation was done with the RNA-seq Kit v2 (Diagenaode), library preparation included 16 cycles of PCR amplification and 150 bp single end reads were sequenced. To obtain a similar amount of *Ca*. E. childressi mRNA reads in each library, we adjusted the amount of reads sequenced per library according to *Ca*. E. childressi mRNA abundance, detailed in **Supplementary information table 16**.

#### LCM transcriptomics: Expression analysis

Expression analysis of LCM RNA reads was done as described in the section “Bulk transcriptomics: Expression analysis” for *Ca*. E. childressi expression, but with additions for the host expression analysis: After removal of non-mRNA contaminants and bacterial contaminants, LCM reads were mapped against the *G. childressi de novo* transcriptome using BBMap (sourceforge.net/projects/bbmap/) at a minimum identity value of 0.85. Mapped reads were counted with FeatureCounts v1.6.3 (Liao et al., 2014), and log-2 fold changes between infection stages were calculated with Aldex2 v3.11 (Fernandes et al., 2013).

### Bioinformatic analyses

#### Genome assembly

All short-read libraries were screened for *Ca*. Endonucleobacter using phyloFlash v3.3 (Gruber-Vodicka et al. 2019). Each short-read library was assembled using Spades v3.7 (Bankevich et al., 2012) after decontamination, quality filtering (trimq = 2) and adapter-trimming of the reads using BBDuk (sourceforge.net/projects/bbmap/). We binned *Ca*. E. childressi and a *Ca*. E. puteoserpentis draft genomes from their respective *G. childressi* and *B. puteoserpentis* metagenomes based on genome coverage, GC content and taxonomic affiliation by using Gbtools v2.6.0 (Seah & Gruber-Vodicka, 2015). *Ca*. E. childressi and *Ca*. E. puteoserpentis short-read genomes were re-assembled by re-mapping the raw short-read libraries to the binned draft genomes by using BBMap (sourceforge.net/projects/bbmap/) with 0.98 minimum identity and Spades v3.7 (maximum k-mer size of 127) (Bankevich et al., 2012). We manually eliminated contigs shorter than 1 kB and screened for contamination using Bandage v0.8.1 (Wick et al., 2015). Next, we calculated quality metrics of both genomes using CheckM v1.0.18 (Parks et al., 2015). For the assembly of *Ca*. E. puteoserpentis high-quality genome, long reads were mapped against *Ca*. E. puteoserpentis short-read genomes using ngmlr v.0.2.7 (Sedlazeck et al., 2018). Output mapped long reads were assembled by using CANU v2.0 (Koren et al., 2017) into *Ca*. E. puteoserpentis long-read genomes. Following an hybrid assembly strategy, the long-read *Ca*. E. puteoserpentis genome was loaded in Unicycler v0.4.8 (Wick et al., 2017) and supplemented with the short-read genome. For the assembly of *Ca*. E. childressi high-quality genome, we assembled PacBio HiFi long reads using CANU. The graphic representation of the resulting long-read metagenomic assembly was loaded into Bandage and *Ca*. E. childressi genome was extracted using *Ca*. E. childressi 16S rRNA as query. We checked *Ca*. E. puteoserpentis and *Ca*. E. childressi high-quality genomes for quality metrics using CheckM v1.0.18 and manually removed contigs shorter than 1 kB. We annotated the genomes using RAST v2.0 (Aziz et al., 2008). We cross-checked RAST annotations manually, and verified the annotations of the main genes discussed in this study by using v.2.10.1 NCBI’s BLAST.

#### Bulk transcriptomics: Expression analysis

We quality trimmed the RNA reads and removed adapters with BBDuk (sourceforge.net/projects/bbmap/). Reads were mapped against RNA and tRNA Silva database v132 (Quast et al., 2012) using BBMap (sourceforge.net/projects/bbmap/) at a minimum identity value of 0.85 to remove non-mRNA contaminants. We pseudoaligned the RNA reads to the hybrid genome of *Ca*. E. childressi using Kallisto v.0.44.0 (Bray et al., 2016) for TPM quantification. Biologically meaningful factors from the global bulk transcriptomic analysis (**Supplementary information table 10)** were used to complement the metabolic reconstruction of *Ca*. E. childressi (**Figure 2, Supplementary information table 3**).

#### Metabolic reconstruction

To model *Ca*. E. childressi metabolism, we loaded its RAST-annotated high-quality genome into Pathway tools v13.0 (Karp et al., 2010). Biologically meaningful factors of pathways representing *Ca*. E. childressi physiology were supplemented with bulk transcriptomic and proteomic data and illustrated in Fig. 2. Pathways for *de novo* synthesis of nucleotides and the bulk transcriptomic expression of all factors involved are covered in Supplementary information table 8. The pathway tools software was also used to assess the completeness of the *de novo* amino acids synthesis capabilities, covered in Supplementary information table 5.

#### Chitinase phylogeny and protein domain analyses

We inferred the function of *Ca*. E. childressi chitinase by determining its phylogenetic affiliation. 38 chitinase amino acid sequences from the G18 glycosidases family were used for this analysis. Sequences included 2 chitinases from *Ca*. E. childressi and *Ca*. E. puteoserpentis, 29 chitinases from diverse *Oceanospirillales* representatives (including *Vibrio cholerae*) and seven chitinases from *Vibrionales* representatives other than *V. cholerae* to root the tree. All sequences were aligned using MAFFT v7.471. The phylogenetic tree was reconstructed using the maximum likelihood-based software IQTREET v1.6.12 using the TIM3 substitution model (1,000 bootstraps). Protein domain analysis of *Ca*. E. childressi chitinase was done using the NCBI online service for protein domain prediction (https://www.ncbi.nlm.nih.gov/Structure/cdd/wrpsb.cgi).

#### *G. childressi de novo* transcriptome: Assembly, curation and annotation

Poly-A RNA reads were quality trimmed and adaptors were removed using BBDuk (sourceforge.net/projects/bbmap/). To remove specific bacterial contaminants, we mapped the reads against the *Ca*. E. childressi and the methane-oxidizing symbiont hybrid genomes. Non-mRNA reads from other potential bacterial contaminants were removed by mapping against the rRNA and tRNA Silva database. We did both mapping steps using BBMap (sourceforge.net/projects/bbmap/) using a minimum identity value of 0.85. After decontamination, we normalized the reads with bbnorm (sourceforge.net/projects/bbmap/), and assembled them with Spades v3.7. We checked the preliminary assembly for completeness and quality metrics using the Trinity Stats package from Trinity v.2.10.0 (Grabherr et al. 2013) and BUSCO v.4.1.2 (metazoan database) (Simão et al., 2015). We assigned taxonomic affiliations to the reads of the preliminary assembly by using blast. Next, we loaded the reads into MEGAN (Huson et al., 2007) and removed non-eukaryotic reads from the preliminary assembly. The resulting assembly was annotated using the trinotate package from Trinity v.2.10.0.

#### IAPs identification

We identified BIR-containing proteins (BIRPs) in *Ca*. E. childressi, *Ca*. E. puteoserpentis and related *Endozoicomonadaceae* genomes by conducting a protein homology analysis. We aligned a total of 48 publically available BIRPs amino acid sequences from tunicates, vertebrates, molluscs, arthropods, entomopoxviruses and ostreid herpesviruses using MAFFT v7.407 (Katoh, 2002; Katoh & Standley, 2013). From this BIRPs alignment we generated a hidden Markov model using the hmmbuild function from hmmer v3.1b2 (Potter et al., 2018) and screened genomes by using the hmmsearch function of hmmer at default thresholds (E-value 1x10E^-3^). Candidate BIRPs were analyzed for functional protein domains using the NCBI online service for protein domain prediction (https://www.ncbi.nlm.nih.gov/Structure/cdd/wrpsb.cgi). We classified those BIRPs having both BIR repeats and RING domains as bona fide inhibitors of apoptosis (**Extended Data Fig. 5**). To verify that the identified IAPs were not mussel host contaminants, we visualized the assembly graph of the *Ca*. E. childressi genome using Bandage v0.8.1 and, using the inbuilt blast function, localized the IAPs within the contigs (**Supplementary Fig. 3**).

#### IAPs phylogeny

Seven amino acid IAP sequences from *Ca*. E. childressi, 13 from *Ca*. E. puteoserpentis, nine from *E. ascidiicola*, two from *E. arenosclerae* and one from *E. numazuensis* were aligned using MAFFT v7.407, together with 20 *G. childressi* host IAP sequences annotated in this study, 17 *B. puteoserpentis* host IAPs annotated in Tietjen, 2020 and 54 publically available IAP sequences from tunicates, vertebrates, molluscs, arthropods, entomopoxviruses and ostreid herpesviruses. The phylogenetic tree was reconstructed using the maximum likelihood-based software IQTREET v1.6.10 (1,000 bootstraps).

#### Phylogenomics and comparative genomics

We analyzed the phylogeny of 514 single-copy genes shared between *Ca*. Endonucleobacter and *Endozoicomonas* representatives. We build a dataset from the *Ca*. E. puteoserpentis and *Ca*. E. childressi genomes generated in this study, 16 publically available *Endozoicomonas* genomes and three *Hahella* genomes as outgroup. Quality metrics of the 21 genomes that formed the dataset are in **Supplementary information table 1**. Prior to the orthology analysis, the *Endozoicomonas* and *Hahella* genomes were annotated using RAST v2.0. We identified the orthologous genes shared among the genomes using OrthoFinder v.2.4.0 (Emms & Kelly, 2019). We identified 514 single-copy genes in the orthology analysis, and extracted a total of 10,794 CDS shared between the dataset genomes. The CDS were aligned using Clustal Omega (Sievers et al., 2011) and the phylogenetic tree was reconstructed using FastTree 2 (Price et al. 2010) (100 bootstraps) as part of the ete3 pipeline from the ETEToolkit (Huerta-Cepas et al. 2016). All genomes were screened for IAPs as described in the “IAPs identification” section above.

#### CRISPR-cas system analysis

We analyzed the *Ca*. E. childressi and *Endozoicomonas ascidiicola*-AVMART05 genomes with CRISPRviz (Nethery & Barrangou, 2019), extracted the nucleotide sequences of the spacers and blasted the spacers against the viruSITE database of complete viral genomes (Stano et al., 2016) by using BLAST+ (Johnson et al., 2008). Blast hits and their P-values are covered in **Supplementary information table 14**. Specific blast hits against herpesviruses are represented in **Supplementary Fig. 1**.

## Acknowledgments

We are grateful to the captains, crew, ROV teams, and chief scientists of the research cruises RV Nautilus NA-58 (2015) and RV Meteor M126 (2016) for their support. We thank Silke Wetzel, Andreas Ellrott and Daniela Tienken from the Max Planck Institute for Marine Microbiology for technical assistance. We thank Bruno Hüttel and Lisa Czaja-Hasse from the Max Planck-Genome-Centre of Cologne for the sequencing and CATS protocol optimization. This study was funded by the Max Planck Society, an ERC Advanced Grant to ND (BathyBiome, 340535), a Gordon and Betty Moore Foundation Marine Microbial Initiative Investigator Award to ND (Grant GBMF3811), the Gottfried Wilhelm Leibniz Prize of the German Research Foundation (DFG) to ND, the USDA National Institute of Food and Agriculture Hatch project 1014212 (MK), and the U.S. National Science Foundation (grants OIA 1934844 and IOS 2003107 to MK). LC-MS/MS measurements were made in the Molecular Education, Technology, and Research Innovation Center (METRIC) at North Carolina State University.

## Author contributions

MAGP, ND and NL conceived the project and designed the experiments. MAGP and NL carried out microscopy experiments. MAGP, AA, HGV and NL performed genome sequencing and analyses. MAGP and AA identified and analyzed IAPs. MAGP and NL designed, optimized and performed the LCM pipeline. MAGP, MT, HGV and NL performed transcriptomes sequencing and analyses. MV and MK performed proteomic data generation and analyses. MAGP, HGV and NL performed phylogenetic analyses. MAGP performed CRISPR and viral analyses. MAGP, ND and NL wrote the manuscript with input from all coauthors.

## Extended Data

**Extended Data Fig. 1.**
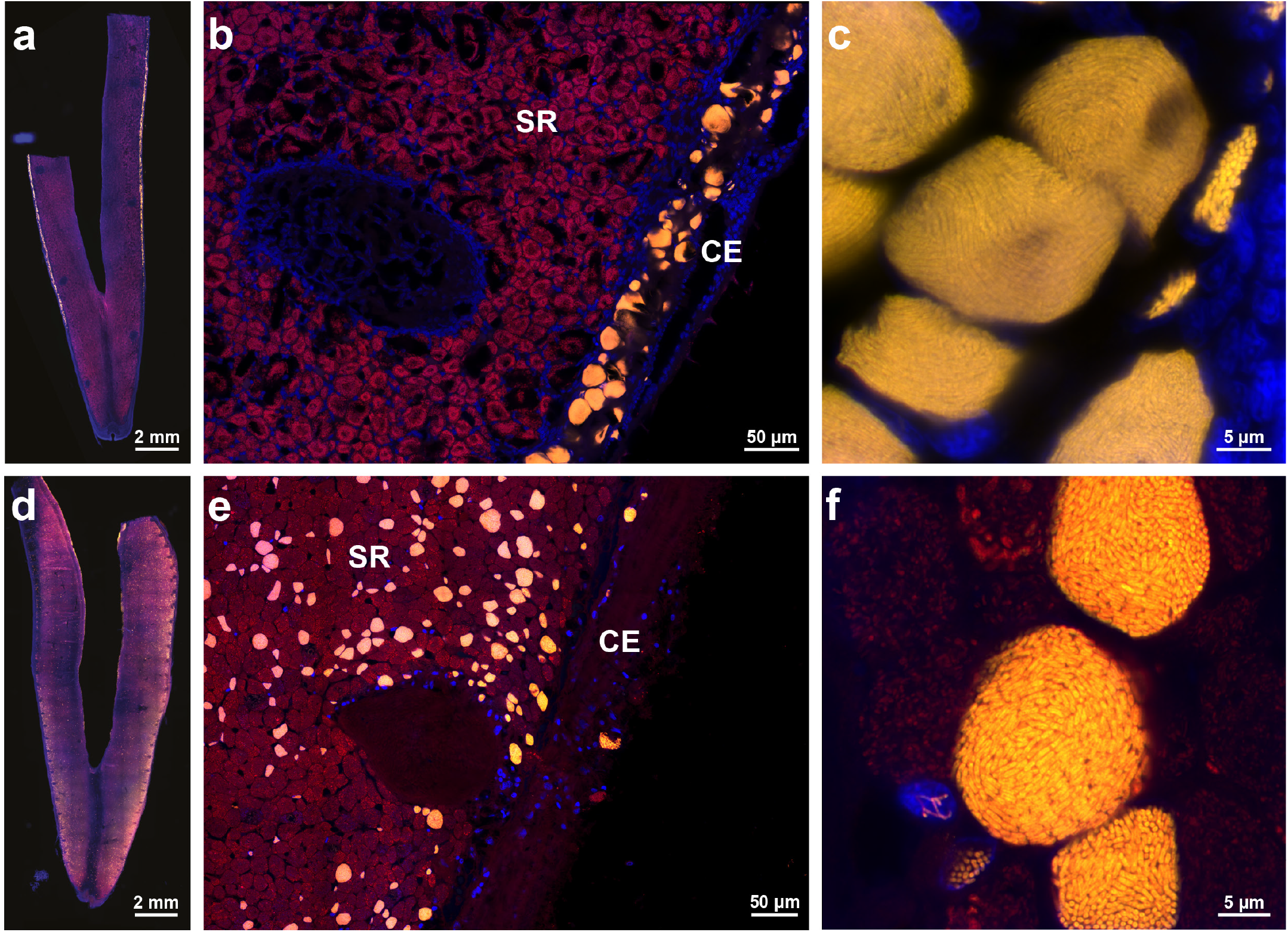
*Ca*. E. childressi only colonized nuclei of cells in the ciliated edges of *G. childressi* gills, while *Ca*. E. puteoserpentis infected the nuclei of cells throughout the gill tissues of *B. puteoserpentis*. ***a*-f**, Fluorescence *in situ* hybridization (FISH) micrographs of single gill filaments of *G. childressi* (**a-c**) and *B. puteoserpentis* specimens (**d-f**) with the FISH probe specific to *Ca*. Endonucleobacter shown in yellow, the eubacterial probe in red, and DAPI-stained DNA in blue. **a** and **d** show stitched overviews of whole gill filaments. **b** and **e** show the symbiotic region (SR) with the sulfur- and methane-oxidizing symbionts, and the ciliated edge (CE), and **c** and **f** show infected nuclei at higher resolution.

**Extended Data Fig. 2.**
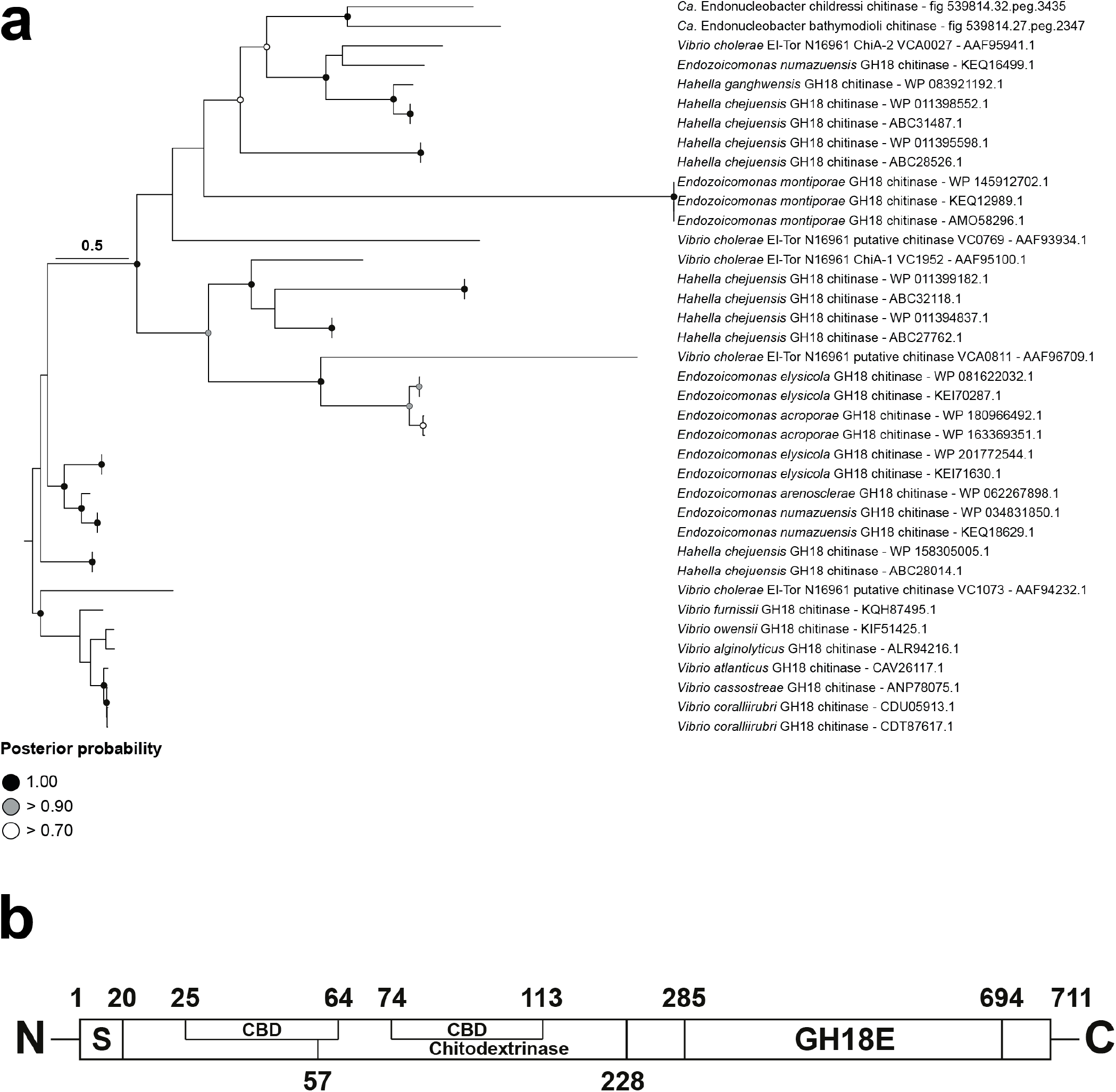
*Ca*. Endonucleobacter chitinases are related to *Vibrio cholerae* chitinase *ChiA-2. Ca*. Endonucleobacter childressi chitinase has a N-terminal peptide for secretion dependent on T3SS. **(a)**, Protein-based phylogeny of 38 chitinases from *Ca*. Endonucleobacter, *Endozoicomonas, Hahella* and *Vibrio* (bootstrap: 1000). We used seven chitinase sequences from *Vibrio* spp. representatives to root the tree. **(b)**, domain analysis of *Ca*. E. childressi chitinase (**S**, T3SS secretion signal peptide; **CBD**, chitin-binding domain; **Chitodextrinase**, chitodextrinase domain; **GH18E**, catalytic domain). Numbers indicate the domain position in the amino acid sequence (not scaled).

**Extended Data Fig. 3.**
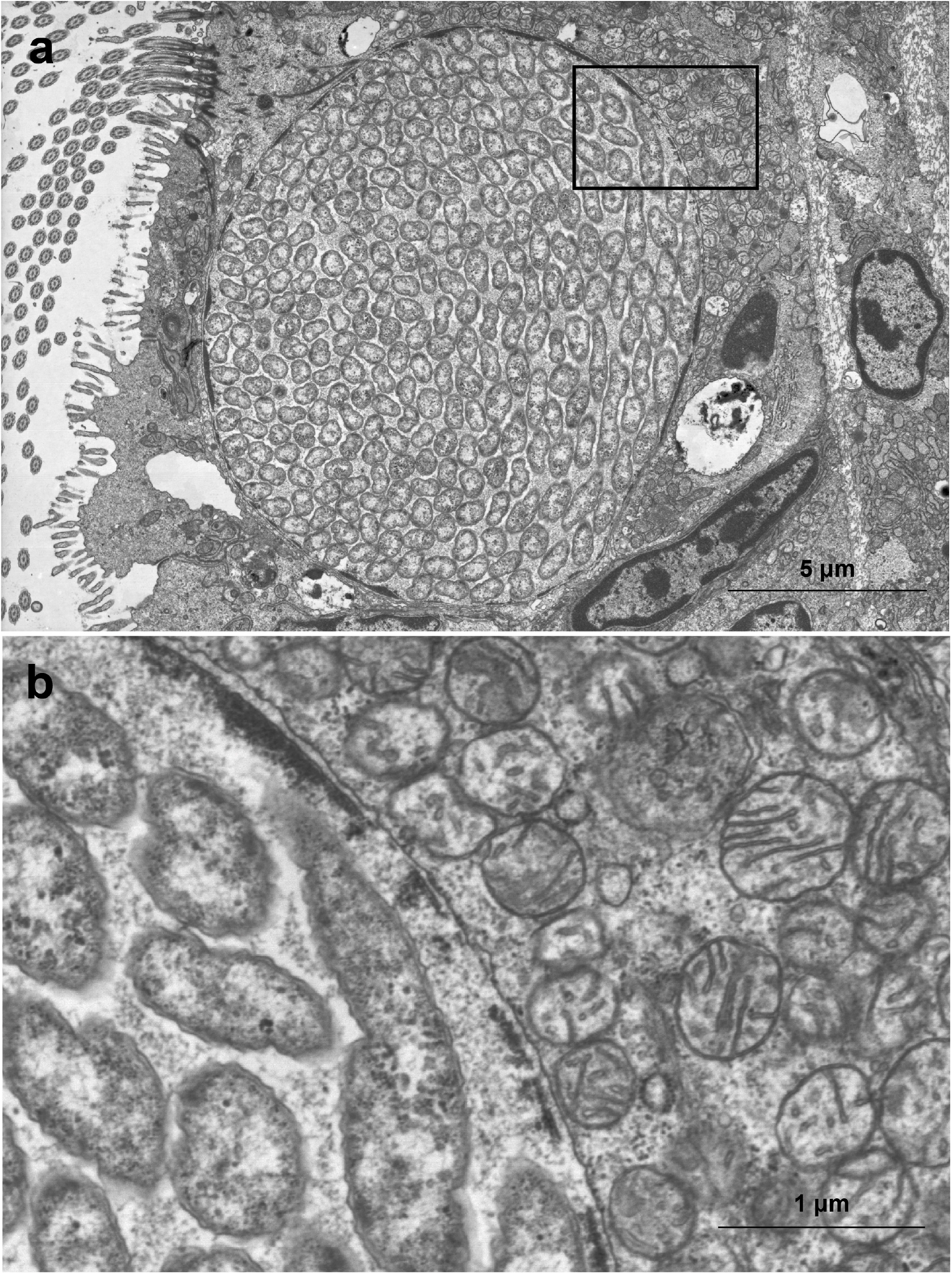
Late infection stage of *Ca*. E. *childressi*. Mitochondria, membranes and other cellular features of the host cell are morphologically intact. **(a)**, TEM overview of a *G. childressi* cell infected by *Ca*. E. childressi. **(b)**, Higher resolution micrograph of rectangle in (a) showing host chromatin compressed along the inner nuclear membrane and morphologically intact mitochondria in the cytosol.

**Extended Data Fig. 4.**
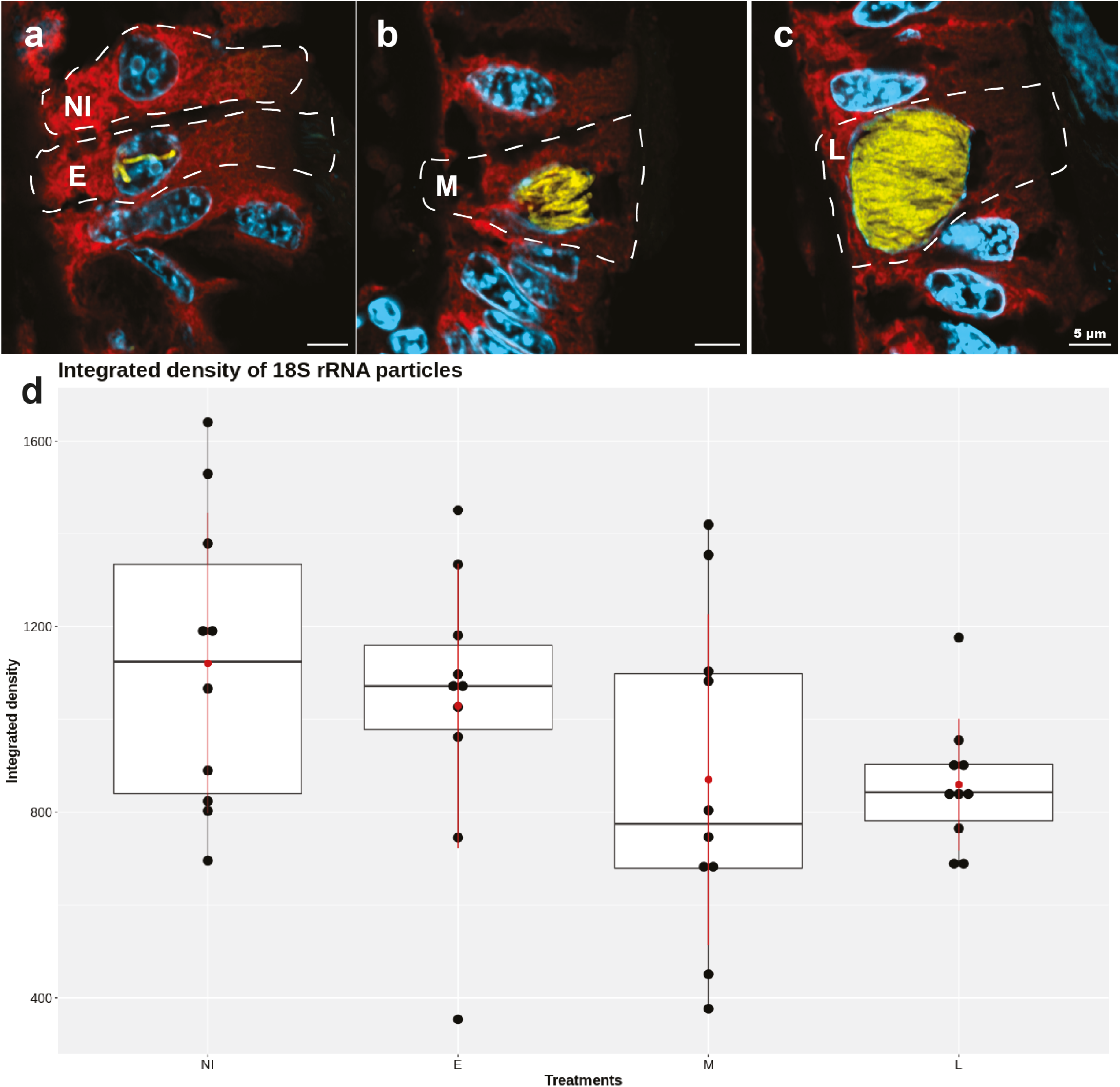
The fluorescent intensity of host 18S rRNA in *G. childressi* cells did not change significantly during the infection cycle. We measured the fluorescent intensity of host 18S rRNA as an indicator of host transcription and ribosomal activity. **(a) - (c):** Fluorescence *in situ* hybridization images of gill cells at different *Ca*. E. childressi infection stages: **(a)** non-infected and early-stage of infection (**NI** = not infected, **E** = early). **(b)** mid-stage of infection (**M** = mide). **(c)** late-stage of infection (**L** = late). **d**, integrated fluorescent intensity of host 18S rRNA normalized to respective cell area.

**Extended Data Fig. 5.**
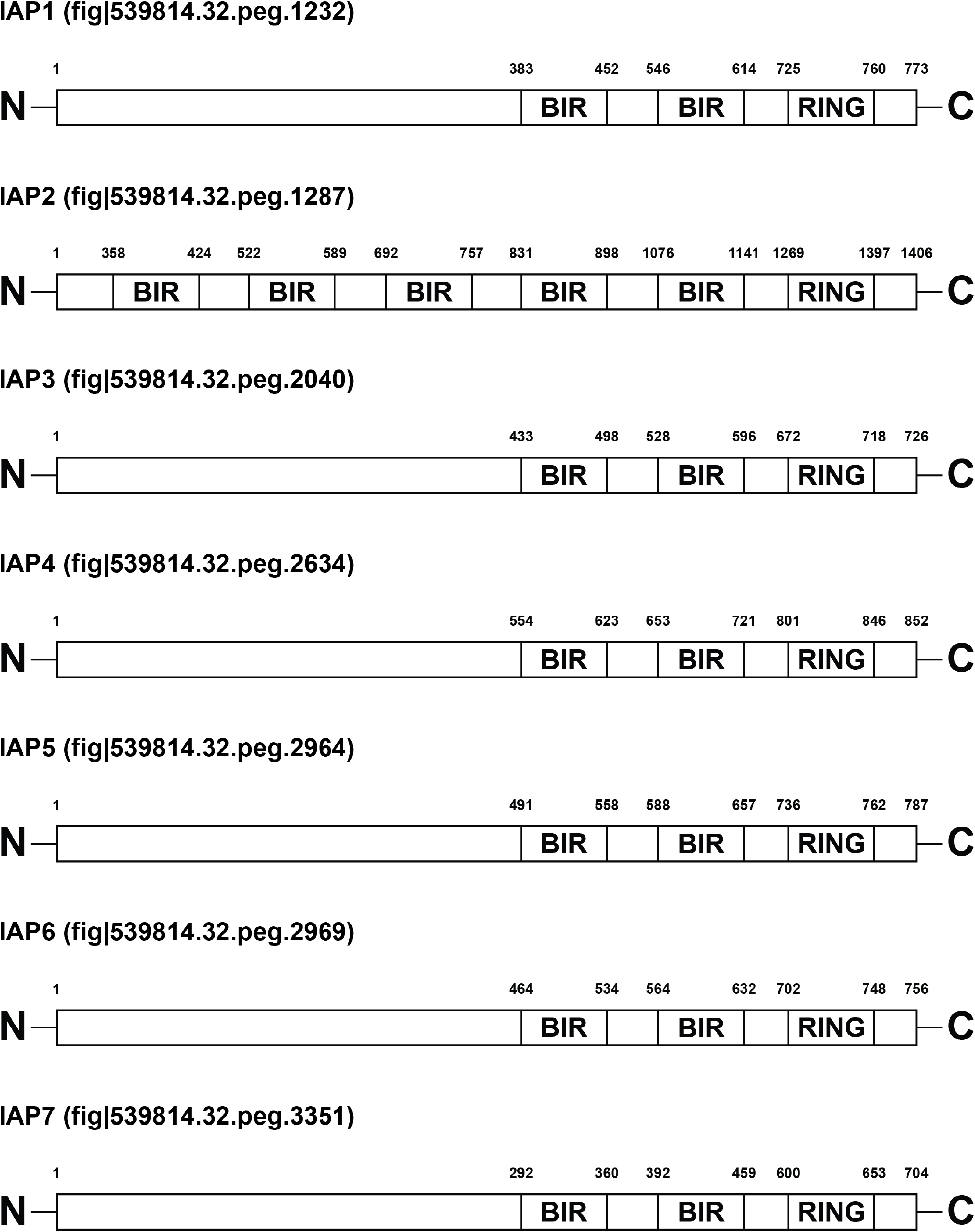
Seven out of the eleven *Ca*. E. childressi BIRPs are bona fide IAPs, with both BIR and RING domains. Domain analysis of the seven IAPs encoded by *Ca*. E. childressi (**BIR**, BIR-repeats domain; **RING**, RING domain). Numbers indicate the domain position in the amino acid sequence (not scaled).

**Extended Data Fig. 6.**
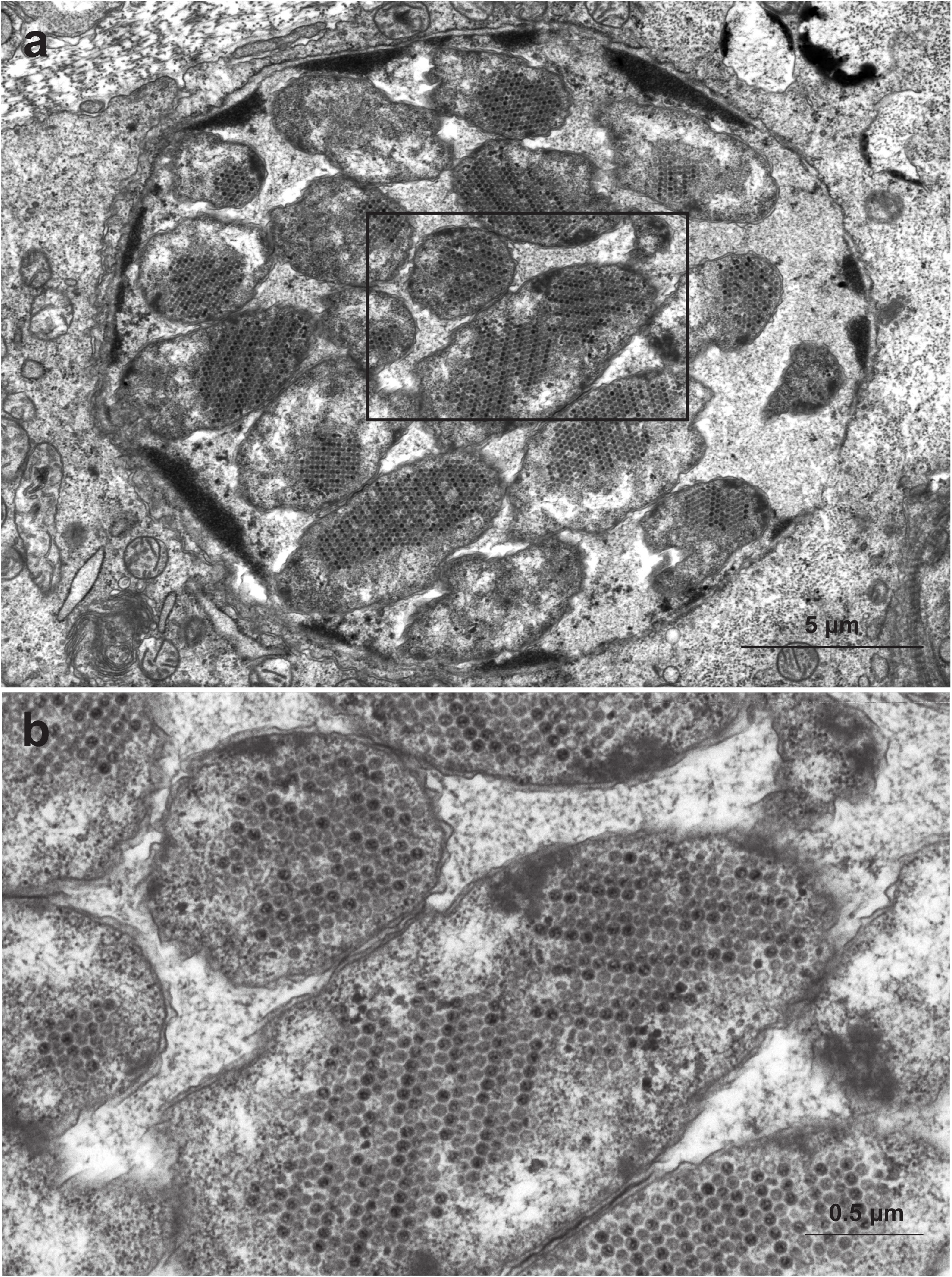
*Ca*. E. childressi inside host nuclei are full of structures typical for viruses. (**a)**, TEM image of *G. childressi* gill cell showing a nucleus infected by Ca. E. childressi that are filled with icosahedral viral capsids. **(b)**, Higher magnification of rectangle in A showing the viral capsids inside Ca. E. childressi cells.

