## Supplementary Information for "An intranuclear bacterial parasite of deep-sea mussels expresses apoptosis inhibitors acquired from its host"

### Contents

**Supplementary note 1 - Description of Ca. Endonucleobacter childressi**

**Supplementary Note 2 – Identification of inhibitors of apoptosis**

**Supplementary note 3 - Ca. Endonucleobacter manipulates host cell cytoskeleton**

**Supplementary note 4 - Nesprin-1 as reporter of nuclear deformation**

**Supplementary note 5 - Possible viral origin of Ca. Endonucleobacter IAPs**

**Supplementary Figure 1-3**

**Supplementary literature**

### Supplementary Notes

#### Supplementary note 1 - Description of *Ca.* Endonucleobacter childressi

Previous work based on the 16S rRNA marker gene, showed high similarities (>98%) between all *Ca*. Endonucleobacter detected in various mussel hosts, and all of them were termed *Ca*. Endonucleobacter bathymodioli. Even though the 16S rRNA is highly similar, the comparison of the average nucleotide identity between the two high quality genome bins resulted in an ANI value of 84.34%, which indicates that these are actually two different bacterial species.

The name "*Ca.* Endonucleobacter childressi" was chosen because of the host species *Gigantidas childressi* in which it can be found. Following a similar line or argument, in this study we decided to rename the originally termed *Ca*. Endonucleobacter bathymodioli described in *Bathymodiolus puteoserpentis* as "*Ca.* Endonucleobacter puteoserpentis" (Zielinski et al., 2009).

The recovered high-quality genome for *Ca.* E. childressi was 3.73 Mb with an average GC of 40.52% and 3,804 protein-coding sequences. The genome bin consists of 12 contigs and has a 98.28 % completeness based on 43 marker genes and no detectable contamination (Supplementary Information Table 1).

The high-quality genome bin of *Ca.* E. puteoserpentis was 3.86 Mb with an average GC of 39.95% and 3,995 protein coding sequences. The genome bin consists of 72 contigs and has a 98.28 % completeness based on 43 marker genes and no detectable contamination (**Supplementary Information Table 1**).

#### Supplementary Note 2 – Identification of inhibitors of apoptosis

An preliminary comparison of bacterial genome annotation pipelines using the JGI pipeline (Nordberg et al., 2014) and the RAST pipeline (Aziz et al., 2008) suggested the presence of IAPs in our bacterial genomes. We therefore generated HMM profiles of 48 publicly available metazoan and viral IAP sequences and performed an hmmsearch (http://hmmer.org/) at default thresholds (E-value $1x10E^{-3}$) against the profile to identify and annotate the IAPs in the bacterial genomes. We differentiated between BIR-containing proteins (BIRPs) and bona fide IAPs in the bacterial genomes by identifying the protein domains of hmmsearch hits by using the NCBI protein domain search platform (https://www.ncbi.nlm.nih.gov/Structure/cdd/wrpsb.cgi). Only those sequences containing a RING domain and at least one BIR-repeat domain were considered as bona fide IAPs (**Extended Data Fig. 5**), while the rest of candidates having a variable number of BIR-repeats but no RING domain were considered as BIRPs. To verify that IAPs were encoded within *Ca*. Endonucleobacter childressi chromosome and not mussel host contamination, we mapped *Ca.* E. childressi IASs within the graphic representation of its genome using Bandage (Wick et al., 2015) (**Supplementary Fig. 3**).

#### Supplementary note 3 - *Ca.* Endonucleobacter manipulates host cell cytoskeleton

As the infection progresses, *Ca.* E. childressi increases the volume of the nucleus up to 50-fold. To understand how *Ca.* Endonucleobacter colonizes and manipulates the host beyond the usage of IAPs, we analyzed its virulence arsenal. Despite its reduced genome, compared to its closest relatives, *Ca*. E. childressi encoded and expressed a broad range of secretion systems. It encoded a T1SS and a T1 dependent RTX adhesin as well as the general secretory (Sec) pathway and the twin-arginine translocation (Tat) pathway, with most genes expressed and the T1 dependent RTX adhesin also present in the proteome (**Supplementary Information Table 10, 11**). Furthermore, *Ca.* E. childressi encoded and expressed the needle-like type 3 secretion system (T3SS), with most components present in the bulk transcriptomic dataset (**Fig. 2**; **Supplementary Information Table 17**).

*Ca.* E. childressi encoded and expressed virulence factors with a T3SS specific N-terminal signal peptide, which are indicative of secretion. Two of these effectors were annotated as the *Shigella*-like factor *IpgD*, and both were present in the bulk transcriptome but absent in the bulk proteome (**Fig. 2; Supplementary Information Table 3, 4**). The enteropathogen *Shigella flexneri* delivers *IpgD*, an inositol phosphate phosphatase, via the T3SS to disentangle the cortical actin of its eukaryotic target and promote cell invasion (Niebuhr et al., 2000). Similarly, *Ca.* E. childressi might be using T3SS-*IpgD* for host cell invasion. However, the expression of *IpgD* and the T3SS translocon factors in the mid and late infection stages suggests that *IpgD* serves an additional function (**Fig. 3; Supplementary Information Table 6**). *IpgD* can interfere with the host cell's actin stress fibers, a molecular corset that provides shape and stiffness to eukaryotic cells (Tavares et al., 2017). By disentangling these stress fibers, *IpgD* can cause cells to lose stiffness and adopt round morphology (Niebuhr et al., 2002). As the infected nucleus massively increases in volume in mid-phase of infection, all eukaryotic organelles will be displaced and pushed against the actin corset. *Ca.* E. childressi might use *IpgD* to reduce mechanical stress and keep its host cell physiologically functional. Overall our analysis shows that *Ca.* E. childressi uses a broad range of secretion systems and effectors to interfere with host cytoskeletal processes. Together with apoptosis inhibition via IAPs, these strategies might work synergistically to ensure the long-term viability of *Ca.* E. childressi replication niche.

#### Supplementary note 4 - Nesprin-1 as reporter of nuclear deformation

To study the host mechanisms that might activate apoptosis, we looked for host factors that were upregulated after *Ca.* Endonucleobacter infection. The host cell upregulated Nesprin-1 in all stages of infection (**Fig. 3; Supplementary Information Table 7**). Nesprins are transmembrane proteins of the nuclear envelope that form part of the nucleoskeleton component of the LINC (linker of nucleoskeleton and cytoskeleton) complex (Bouzid et al., 2019). Within the LINC complex, nesprin-1 plays a key role as a biomechanical reporter of nuclear position and morphology (Zhang et al., 2010). Infected *G. childressi* cells might react to *Ca.* E. childressi growth-induced nuclear expansion by upregulating nesprin-1, in an attempt to compensate irregularities in nuclear position and morphology. In model organisms, alterations of the actin cytoskeleton of the cell are common triggers of apoptosis (Kräter et al., 2018; White et al., 2001) (reviewed in Desouza et al., 2012). Intriguingly, the highest expression of IAPs by *Ca.* E. childressi occurred in mid-stage of infection, when *Ca*. Endonucleobacter will cause nuclear expansion and via Nesprin-1 transfer this deformation to the host cytoskeleton (**Fig. 3; Supplementary Information Table 6**).

#### Supplementary note 5 - Possible viral origin of *Ca.* Endonucleobacter IAPs

The ostreid herpes virus OsHV-1 (Herpesvirales, Mallacoherpesviridae) is a virus of molluscs that encodes for IAPs (Davison et al., 2005). Unlike other herpesviruses, OsHV-1 is not restricted to a single host species, infecting oysters, clams, scallops and octopuses (Arzul et al., 2001a; Arzul et al., 2001b; Hine et al., 1998; Hine & Thorne, 1997; Prado-Alvarez et al., 2021). To investigate if ostreid herpesviruses mediated HGT of IAPs from mussels to *Ca.* Endonucleobacter, we included OsHV-1 IAPs sequences in our phylogenetic study (**Fig. 4**). Interestingly, both OsHV-1 IAP were interspersed with those of *Ca.* Endonucleobacter, *Endozoicomonas*, molluscs and ascidians (**Fig. 4**). We looked for evidences of viral-mediated HGT of BIRPs and IAPs from *G. childressi* to *Ca.* E. childressi. *Ca.* E. childressi is subjected to viral predation while sheltered in the nucleus of its host cell (**Extended**

**Data Fig. 6**). We could recover around 17% of the OsHV-1 genome from the same *G. childressi* specimen from which we assembled the *Ca.* E. childressi genome. This indicated that *Ca.* E. childressi and OsHV-1 coexist in the same mussel host. Some bacteria have an adaptive immune system called CRISPR-cas, which prevents viral predation by integrating viral sequences called spacers (Rath et al., 2015). Three spacers from the CRISPR-cas systems 1 and 3 of *Ca.* E. childressi showed homology to *Herpesvirales* (**Supplementary Fig. 2; Supplementary Information Table 14**). Viruses have been traditionally classified into archeoviruses, bacterioviruses and eukaryoviruses based on the life domain of their host (Krupovic et al., 2016). Although host jumps within life domains are frequent, viruses are not known to simultaneously infect hosts from different life domains (Geoghegan et al., 2017; Longdon et al., 2014). Given their ecology, viruses that infect and/or persist as integrated prophages of bacterial symbionts of eukaryotic cells are simultaneously exposed to different life domains (Brüssow et al., 2004; Turnbaugh et al., 2007). Importantly, such interactions blur the traditional concept of "virus host" and raise the possibility of viruses interacting (not necessarily in a lytic manner) and exchanging genetic material simultaneously with more than one superkingdom of life (Malik et al., 2017). It is tempting to speculate that OsHV-1 or a yet to be discovered close relative might have mediated HGT of BIRPs and IAPs from animal hosts to Endozoicomonadaceae.

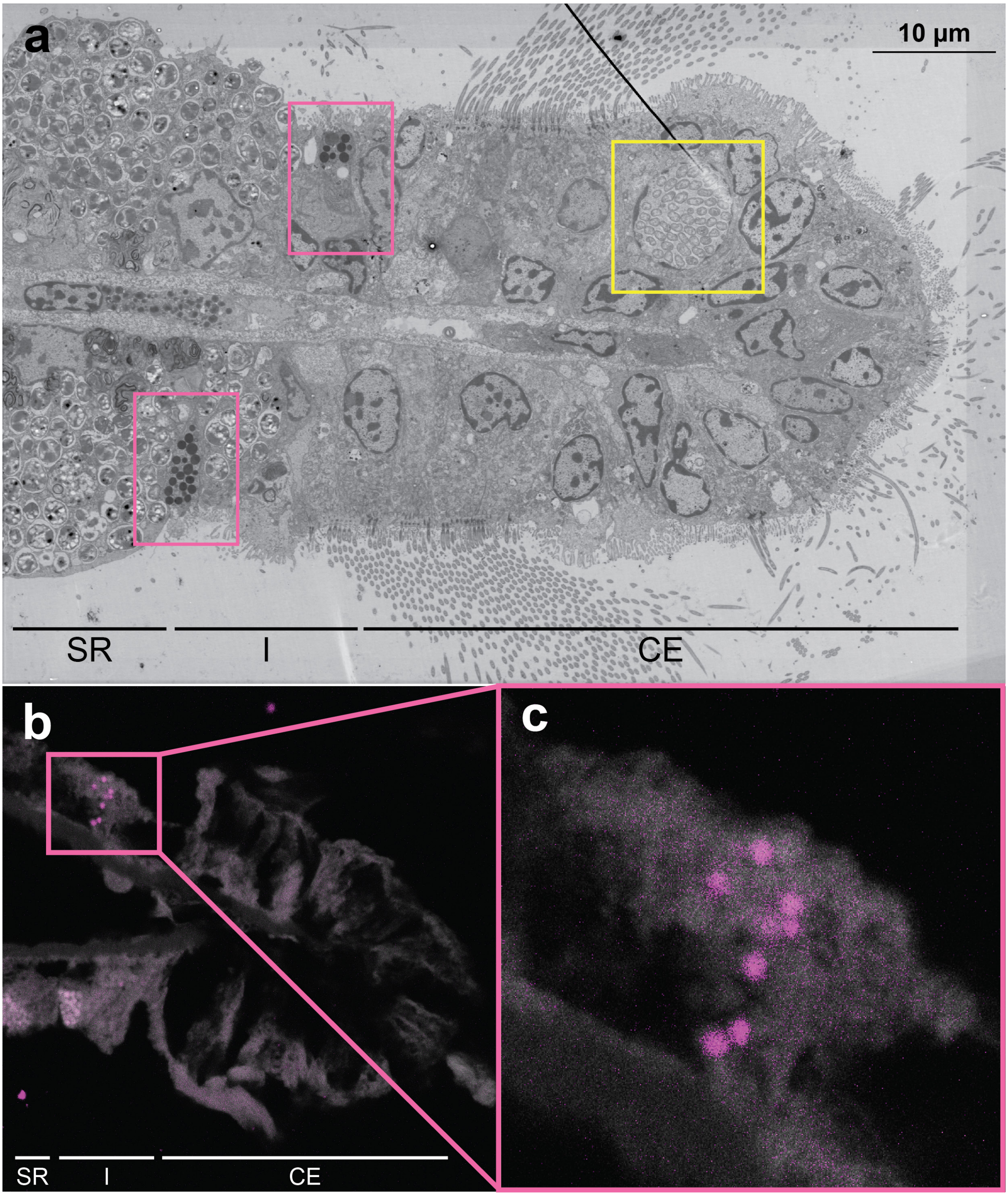


**Supplementary figure 1 | Non-infected *G. childressi* secretory gill cells secrete mucins to the extracellular medium. (a)**, TEM overview of cross section of *G. childressi* gill filament (**SR**: Symbiotic region, **I**: Interface, **CE**: Ciliated edge). Pink frames highlight secretory cells, while the yellow frame highlights a ciliated edge cell infected by *Ca.* E. childressi. **(b)**, Airyscan overview of cross section of *G. childressi* gill filament. N-acetylglucosamine residues (pink: FungiFluor™), *G. childressi* gill tissue (grey: Autofluorescence). **(c),** amplification of pink frame in **b** highlighting the polarity of the secretory cell and the directionality of the vesicles.

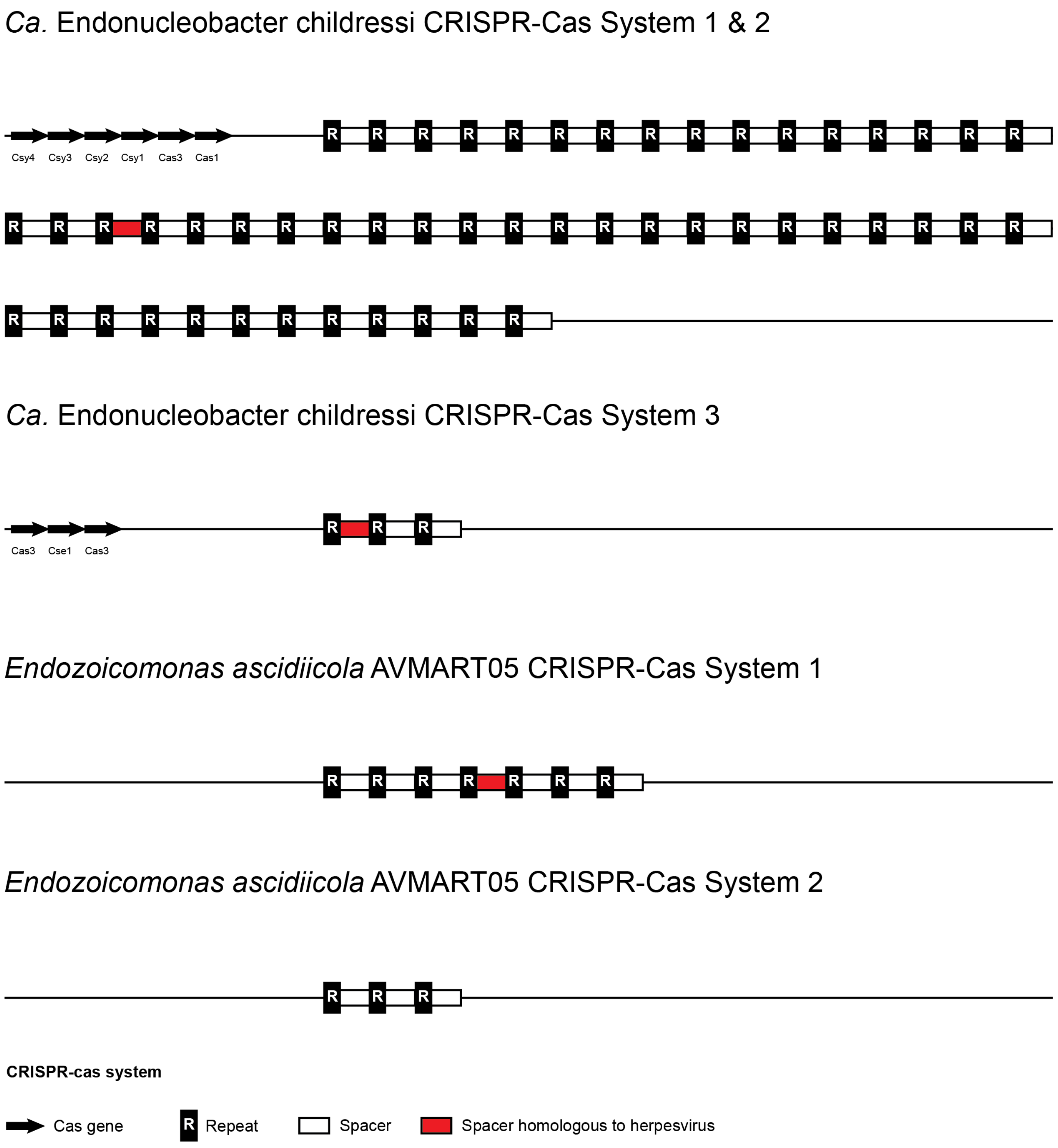


**Supplementary figure 2 | *Ca.* E. childressi and *E. ascidiicola* CRISPR spacers have high similarity to herpes virus genomes.** The *Ca.* E. childressi CRISPR array had two spacers with high similarity to the genomes of herpes viruses, while *E. ascidiicola* had one spacer with high similarity to herpes virus genomes.

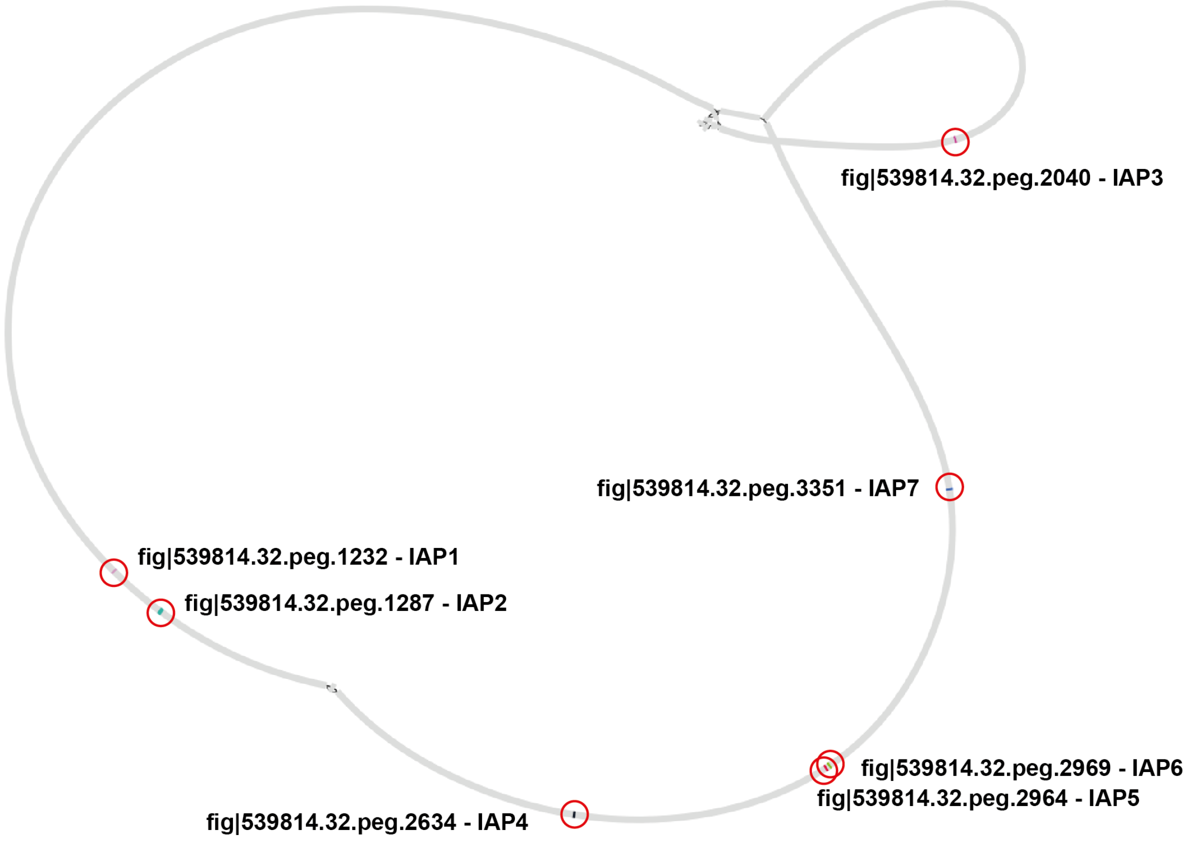


**Supplementary figure 3 | All seven IAPs encoded by *Ca.* E. childressi are located on the main contigs of its chromosome.** Graphic representation of *Ca.* E. childressi genome assembly (Bandage). Red circles point the exact location of IAPs within their respective contigs.
